# diploS/HIC: an updated approach to classifying selective sweeps

**DOI:** 10.1101/267229

**Authors:** Andrew D. Kern, Daniel R. Schrider

## Abstract

Identifying selective sweeps in populations that have complex demographic histories remains a difficult problem in population genetics. We previously introduced a supervised machine learning approach, `S/HIC`, for finding both hard and soft selective sweeps in genomes on the basis of patterns of genetic variation surrounding a window of the genome. While `S/HIC` was shown to be both powerful and precise, the utility of `S/HIC` was limited by the use of phased genomic data as input. In this report we describe a deep learning variant of our method, `diploS/HIC`, that uses unphased genotypes to accurately classify genomic windows. `diploS/HIC` is shown to be quite powerful even at moderate to small sample sizes

## Introduction

An important goal of population genetics is to accurately identify loci in the genome which have undergone recent selective sweeps. In natural populations with complex demographic histories this is a difficult proposition, and thus much attention has been devoted to improving sweep-finding methods (e.g. Nielsen et al. [2005], Lin et al. [2011], DeGiorgio et al. [2016]). The vast majority of this work has centered upon a “hard” sweep model of adaptation, wherein a *de-novo* beneficial mutation sweeps to high frequency carrying with it its linked genetic background [Maynard Smith and Haigh, 1974, Kaplan et al., 1989]. While such hard sweeps may be important sources of adaptive evolution, there is good reason to believe that selection from standing variation, a.k.a. “soft” sweeps, might be equally if not more important. Indeed, both theoretical and empirical evidence have accumulated sufficiently as to suggest that soft sweeps might be the more frequent mode of adaptation in many natural populations (Hermisson and Pennings [2005], Messer and Petrov [2013], Garud et al. [2015], Sheehan and Song [2016], Schrider and Kern [2017] but see Jensen [2014]). Thus if we wish to generate a more complete catalog of the genomic targets of selection, we need to be able to reliably identify both hard and soft selective sweeps.

While this sounds straightforward at first blush, a lingering issue is that hard sweeps can generate spurious signatures of soft and partial sweeps in regions that are at intermediate genetic distances from the true target of selection (Schrider et al. [2015]). This “soft-shoulder” effect thus requires any method developed for finding soft or partial sweeps to be aware of the genomic spatial context. We recently developed a method known as `S/HIC` that meets this criterion and is able to find both hard and soft sweeps in the genome with unparalleled accuracy and robustness (Schrider and Kern [2016]). `S/HIC`, like several contemporary tools for sweep finding (e.g. Pavlidis et al. [2010], Lin et al. [2011], Ronen et al. [2013], Pybus et al. [2015], Sheehan and Song [2016]), utilizes supervised machine learning to classify individual windows of the genome as sweeps. In particular `S/HIC` summarizes a genomic window on the basis of a large vector of transformed summary statistics and then uses those “features” as input to an Extra-Trees classifier (Geurts et al. [2006]). The feature vector of `S/HIC` captures spatial patterns of variation across a region of the genome by recording the relative values of a given statistic in subwindows as a vector. While `S/HIC` performs well both on simulated and empirical data, its use is currently limited to population genomic samples with phased genotype data as it relies on haplotype-based statistics.

Here we introduce a new version of `S/HIC` that alleviates this issue that we call `diploS/HIC` (pronounced “deep-lo-shick”). We outline a series of summary statistics which we calculate on unphased genotypes across a genomic window which allow for accurate classification of genomic windows in the context of supervised machine learning (c.f. Schrider and Kern [2018]). `diploS/HIC` uses a deep convolutional neural network (CNN) approach to classification (LeCun et al. [2004], Krizhevsky and Hinton [2010], Krizhevsky et al. [2012]) whereby we represent as an image the multidimensional vector of statistics calculated from the window to be classified. Using simulation we show that `diploS/HIC` retains nearly all of the power and accuracy of `S/HIC`. Moreover we show that convolutional neural networks outperform our original Extra-Trees classifier both for our original `S/HIC` formulation and `diploS/HIC`.

## Methods and Model

### Coalescent simulation

`diploS/HIC` is a supervised machine learning classifier that uses coalescent simulation to generate training data. All of our simulations in this paper are performed using discoal (Kern and Schrider [2016]), though other simulators could be used provided they are flexible enough to simulate the evolutionary scenarios described below. We simulated under a number of demographic scenarios and selective scenarios both with constant population size and with the very strong population growth we observed in *Anopheles gambiae* from Burkina Faso (Anopheles gambiae 1000 Genomes Consortium et al. [2017]). These simulations were performed under five sample sizes, *n* = *{*20, 40, 60, 80, 100*}* hap-loid chromosomes. Command lines for each scenario are given in Supplemental Table 1. For simulations with selective sweeps we sampled from a uniform prior of population scaled selection coefficients (*α* = 2*Ns*). For each combination of sample size, demographic history, and rage of *α*, we simulated large chromosomal regions that we later split into 11 subwindows (as in Schrider and Kern [2016]). We simulate hard sweep training examples by generating 2 *×* 10^3^ simulations where we conditioned upon a fixation in the exact center of our region (i.e. the middle of the 6^th^ subwindow). Linked-hard training examples are generated by simulating 2 *×* 10^3^ sweeps where the selected site is at the center of each of the remaining 10 subwindows flanking the central window. For soft sweep and linked-soft training examples we follow this same procedure but add a uniform prior on the frequency, *f*, at which a mutation is segregating at the time it becomes beneficial such that *f ∼ U* (0.0, 0.2). Most simulations under constant population sized varied the time of fixation of the beneficial allele, *τ*, over a uniform range *τ ∼ U* (0, 0.025) in units of 4*N* generations, although where noted we fix *τ* = 0. Our simulations using *Anopheles gambiae* demography were done with a more recent distribution of fixations times, *τ ∼ U* (0, 0.0004). Finally we generate 2 *×* 10^3^ simulations without sweeps but with the same demographic history as selected training examples.

From these simulations a balanced set with equal representation of all five classes (neutral loci, hard sweeps, soft sweeps, loci linked to hard sweeps, and loci linked to soft sweeps) to be used for machine learning is created through sampling without replacement. This constitutes the training set for our classifier. The simulation procedure is then repeated to generate an independent test set of’ 10^3^ examples per class on which to benchmark trained classifiers.

### The **diploS/HIC** feature vector and associated summary statistics

Our original version of `S/HIC` used as a feature vector the transformed values of 11 summary statistics calculated in 11 sub-windows centered upon the window to be classified (Schrider and Kern [2016]). These statistics were *π* (Tajima [1983]), *θ*^^^*_w_* (Watterson [1975]), Tajima’s *D* (Tajima [1989]), *θ*^^^*_H_* and Fay and Wu’s *H* (Fay and Wu [2000]), the number of distinct haplotypes, average haplotype homozygosity, *H*_12_ and *H*_2_*/H*_1_ (Garud et al. [2015]), *Z_ns_* (Kelly [1997]), and the maximum value of *ω* (Kim and Nielsen [2004]). These statistics are then normalized across subwindows to capture the relative shape of a given statistic across the larger region such that the value of some statistic *x* in the *i*th subwindow is *x_i_* = *x_i_/ _j_ x_j_*. If the minimum value of *x* is negative, the elements of the vector are rescaled such that *x_i_* = *x_i_ − min*(*x*). Classification of genomic windows then proceeds using an Extra-Trees classifier (ETC), and windows were identified as belonging to one of five classes: a hard sweep, a soft sweep, a window linked to a hard sweep, a window linked to a soft sweep, or a “neutral” window unlinked to a sweep. Given our choice of summaries, `S/HIC` relies upon phased haplotype data (e.g. *H*_12_, *Z_ns_*) and a polarized site frequency spectrum (SFS) (Fay and Wu’s *H* and *θ*^^^*_H_*), though the latter can be removed with little loss of power (data not shown).

To relax these constraints we set out to design a feature vector that would allow for genomic windows to be classified into our five classes on the basis of a set of statistics that could be calculated from unphased genotype data. In particular we aim to capture quantities that describe three axes of the data: the SFS, haplotype structure in the region (we will only have partial information about this), and linkage disequilibrium (LD).

Unphased data do not lead to difficulties in summarizing the SFS, so we use three summary statistics: *π* (Tajima [1983]), *θ*^^^*_w_* (Watterson [1975]), and Tajima’s *D* (Tajima [1989]). Describing the haplotype structure and LD from unphased data is more difficult. Let the genotype vector of the *k*th individual be *x_k_* = *{x_k_*_0_*, x_k_*_1_*, …, x_km_}* where individual *k* has been genotyped at *m* loci and *x_ki_ ∈ {*0, 1, 2*}* according to the number of non-reference (minor) alleles individual *k* has at locus *i*. We define the genotype distance, *g_kl_*, between two individuals *k* and *l* such that 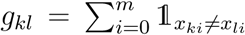. We note that *g_kl_* is an underestimate of the true sequence divergence, and thus only provides a lower bound on haplotype distance. While that is so, moments of the distribution of pairwise *g_kl_* comparisons for a sample prove informative for differentiating among our sweep and neutral classes of loci. In particular we use the variance, the skewness, and the kurtosis of the *g_kl_* distribution as summary statistics.

Figure 1 shows mean values of the normalized subwindow *g_kl_* distribution moments from constant size population simulations of hard and soft sweeps (see figure legend for details). While each of these moments contains information about the location of a sweep, the kurtosis in particular adds information as to whether a given sweep is soft vs. hard. Here we are using this for diploid genotype data, however we note that a similar summary of haploid data (i.e. the higher order moments of haplotype distances) might add excellent information for sweep classification from phased data.

**Figure 1:**
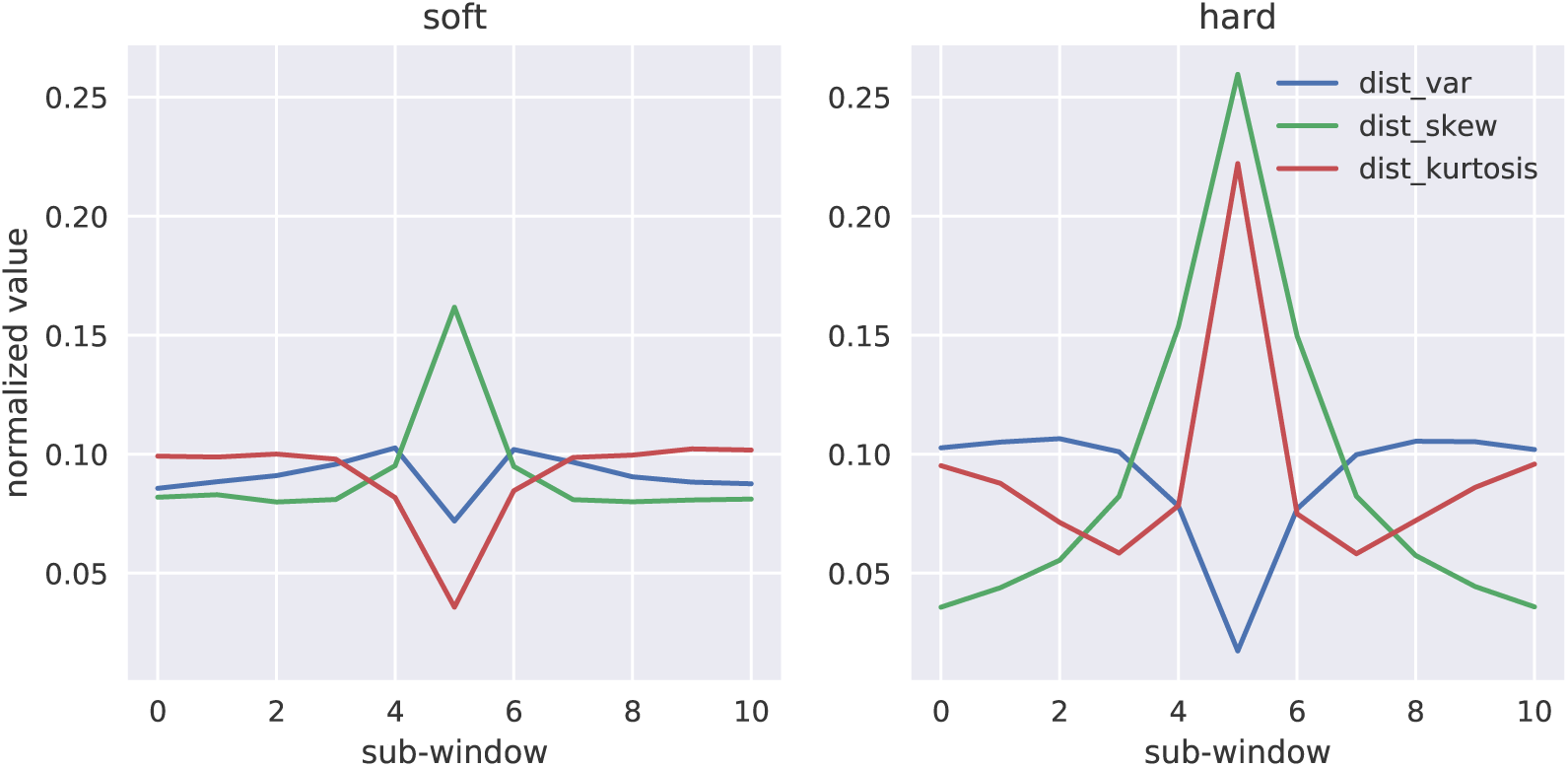
Genotype distance distribution summaries for soft and hard sweeps. We simulated 2 10^3^ sweeps under constant population size with moderately strong selection (*α ~ U* (250, 2500)), in which the fixation of the beneficial allele happened immediately prior to sampling (i.e. *τ* = 0). The relative magnitude of the variance, skew, and kurtosis of the distribution of pairwise genotype distances, *g_kl_*, across sub-windows of a larger simulated region is shown for soft and hard sweeps respectively

We also summarized the frequency distribution of multilocus genotypes in a manner analogous to the haplotype frequency summary statistics of Messer and Petrov [2013] and Garud et al. [2015]. The simple idea here is to treat the multilocus genotype as a population genetic entity, sometimes referred to as the “diplotype”. We recorded four summaries: the number of distinct genotype vectors, and then the multilocus genotype equivalents of *H*_1_, *H*_12_, and *H*_2_*/H*_1_, call them *J*_1_, *J*_12_, and *J*_2_*/J*_1_. The normalized values of *J*_1_, *J*_12_, and *J*_2_*/J*_1_ across simulated genomic regions that contain either a soft or hard selective sweep are shown in Figure 2. Each of these statistics adds useful information for detecting hard sweeps, however they are less sensitive to soft sweeps than the *g_kl_* distribution introduced above. Thus, these statistics may be useful for discriminating between hard and soft sweeps. Finally, to summarize patterns of linkage disequilibrium we computed the equivalents of *Z_ns_* (Kelly [1997]) and *ω* (Kim and Nielsen [2004]) using genotypic LD rather than gametic LD calculated according to Rogers and Huff [2009].

**Figure 2:**
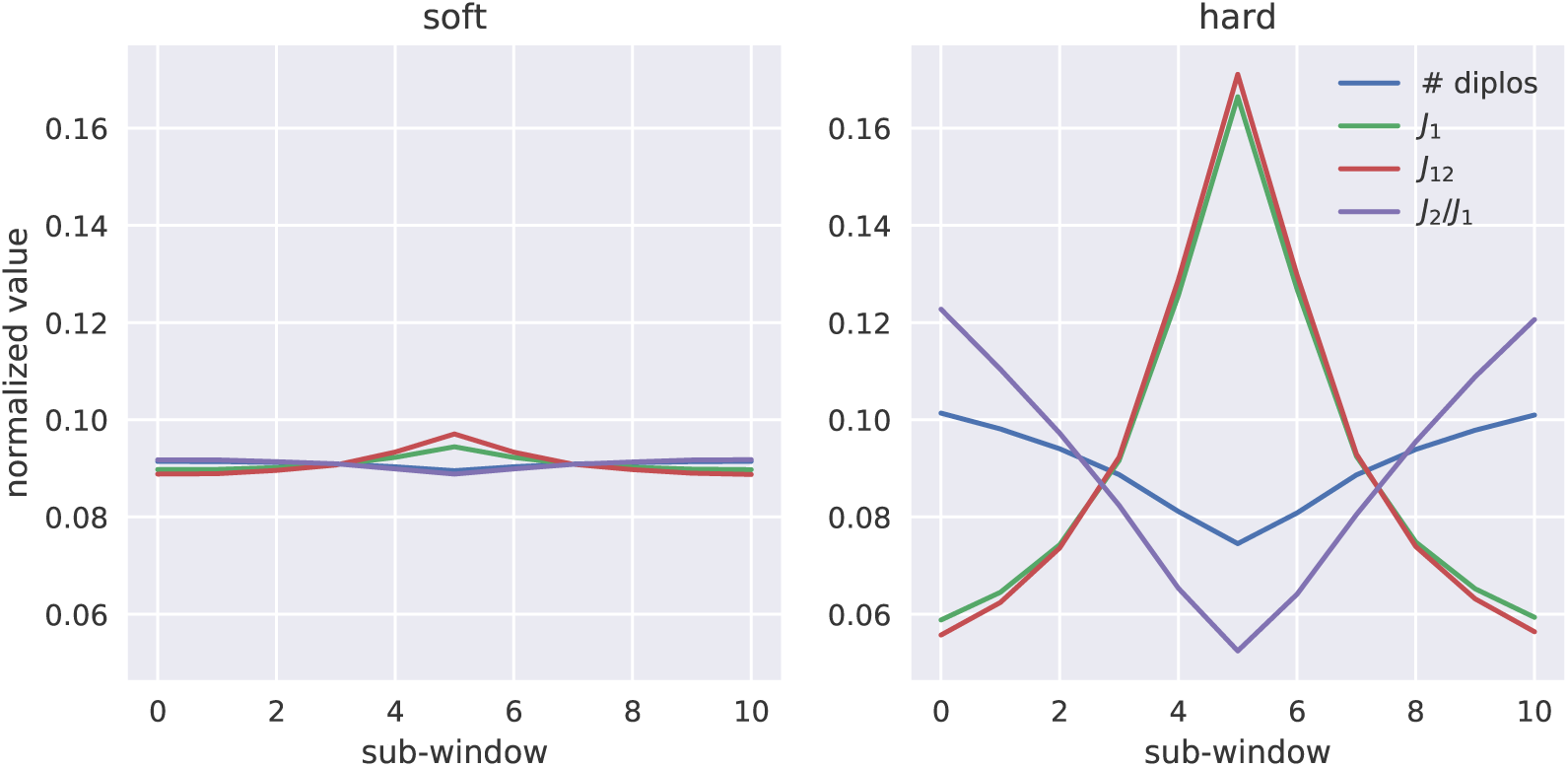
Multilocus genotype frequency spectrum statistics for soft and hard sweeps. We simulated 2 10^3^ sweeps under constant population size with moderately strong selection (*α ~ U* (250, 2500)) and in which the fixation of the beneficial allele happened immediately prior to sampling (i.e. *τ* = 0). The relative magnitude of the number of diploid genotypes, *J*_1_, *J*_12_, and *J*_2_*/J*_1_ across sub-windows of a larger simulated region is shown for soft and hard sweeps, respectively

Thus our complete feature vector for `diploS/HIC` consists of 12 summary statistics calculated in each of our 11 sub-windows (132 in total): *π*, *θ*^^^*_w_*, and Tajima’s *D*, the variance, the skew, and the kurtosis of *g_kl_*, the number of multilocus genotypes, *J*_1_, *J*_12_, *J*_2_*/J*_1_, unphased *Z_ns_*, and the maximum value of unphased *ω*.

### Sweep finding as image recognition

In our original implementation of `S/HIC` we used an Extra-Trees classifier to classify genomic windows on the basis of a large vector of summary statistics that had been transformed to capture spatial information. While the classifier was thus able to use spatial information, the actual implementation was completely naive to any notion of spatial structuring among subwindows. In an effort to utilize spatial relationships of summary statistics among windows more effectively, we reasoned that we might be able to represent our transformed summary statistics as an image that we could then train a convolutional neural network (CNN) to recognize. As before, we record our vector of 12 summary statistics across 11 subwindows in which our central window is the one to be classified. Putting summary statistics on rows and subwindows on columns we then arrive at an image representation of a locus on the basis of our collection of summary statistics (Figure 3). In Figure 3 we present averages across each class and in Supplemental Figure S1 we show examples of several individual simulation images; note that any given individual realization shown in Supplemental Figure S1 can differ substantially from the mean. These are the raw material that are used for training our CNN classifier.

**Figure 3:**
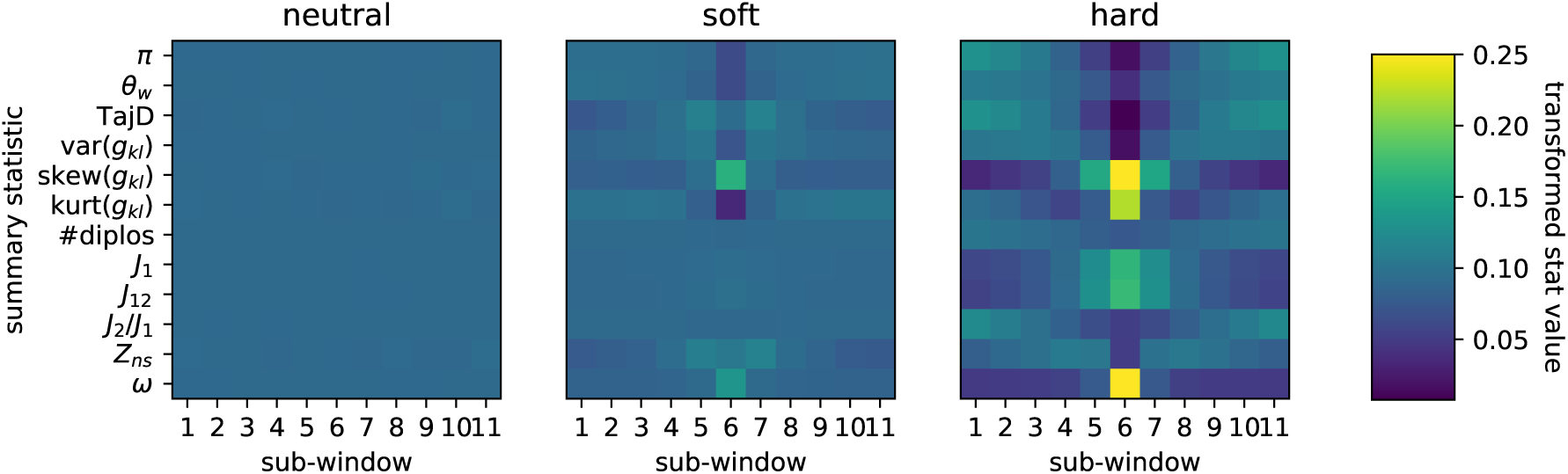
Average-case images for neutral regions, soft sweeps, and hard sweeps. Here we show the average images for three classes of training data, neutral regions, soft sweeps, and hard sweeps, that are used as input to our convolutional neural network for classification. Individual subwindows of the region to be classified are shown in columns. Rows represent each of the 12 summary statistics (see text). The upper end of the color scale has been truncated at 0.25 for visualization. These two images represent the average image from a set of 2 10^3^ simulations with parameter values as described in Figure 1.

### Convolutional Neural Network

To classify genomic regions into one of our five classes (a hard sweep, a soft sweep, a window linked to a hard sweep, a window linked to a soft sweep, or a “neutral” window unlinked to a sweep) `diploS/HIC` uses a deep convolutional neural network (CNN) trained from simulation data. CNNs are characterized by having one or more “convolutional” layers in their networks whose purpose is to capture essential features of an image by sliding a receptive field of specified size across the image an then computing dot products between the original image values and that of the convolutional filter. CNNs are incredibly powerful tools for image recognition because they reduce the number of parameters necessary for deep learning on images while maintaining enough of the complexity of the original data for accurate classification or regression (LeCun et al. [2004], Krizhevsky and Hinton [2010], Krizhevsky et al. [2012]).

`diploS/HIC` uses a CNN architecture that attempts to capture relationships between windows at multiple physical scales. To do this our input image is used as input to three different branches of a CNN, each of which has two dimensional convolution layers with different sized filters (see Figure 4). Each of these convolutional layers slides a “receptive filter,” basically a focal window, across the input image, taking dot-products between filter weights (i.e. the fitted parameters of the network layer) and the input itself. Doing this at multiple scales simultaneously allows us to capture how summary stats are responding across the physical chromosome more effectively. The top-most “convolutional unit” has two 3 *×* 3 convolutional layers, where as the other two convolutional units have 2 *×* 2 filters that are dilated such that the receptive filter is rectangular–the middle unit uses a 1 *×* 3 dilation and the bottom unit uses a 1 *×* 4 dilation (*c.f.* Yu and Koltun [2015]). In experiments that we do not show here, this structure was found to outperform simpler CNNs that did not attempt to capture multiscale information. Moreover, while the two dimensional convolution structure implies that the summary statistic row ordering might matter, we experimented with different permutations and observed no discernible effect on accuracy. Each of these convolutional units consists of two convolutional layers with ReLu activations, followed by a max pooling layer (to reduce the dimensionality of the representation), which itself is followed by a dropout layer to control overfitting by regularizing the network through setting a subset of weights to zero stochastically. This combination of convolutional layers followed by pooling and dropout layers is quite standard in image recognition (see O’Shea and Nash [2015] for an approachable introduction). At this point in the network the outputs from all three convolutional units are flattened and concatenated and then fed to a series of two fully connected dense layers, between which we again utilize dropout layers, before finally arriving at a softmax activation layer that outputs categorical class membership probabilities (Figure 4).

**Figure 4:**
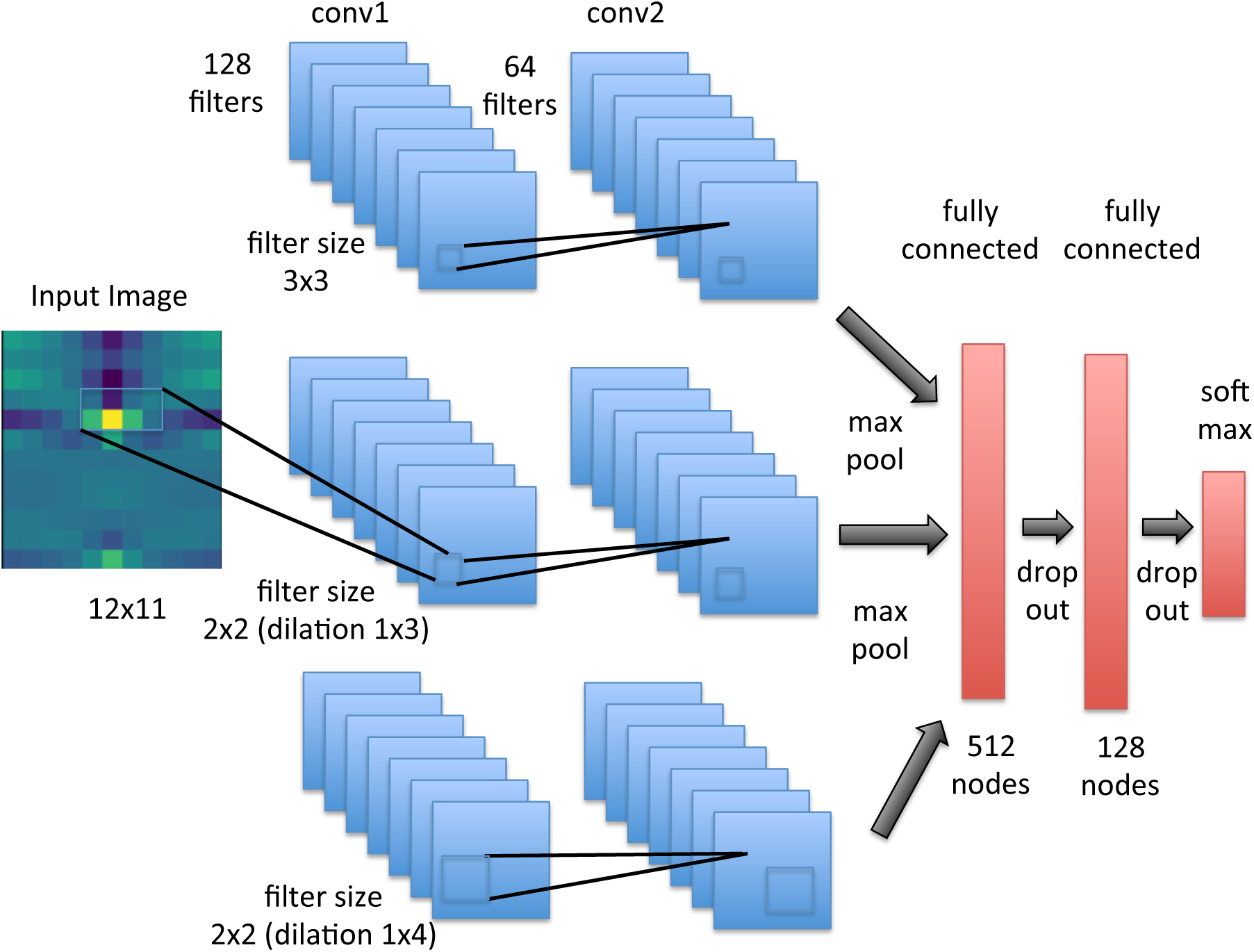
Convolutional neural network structure of diploS/HIC. The CNN takes a single image as input and then passes that image to three three convolution layer units each with different filter sizes to capture variation in the image at different physical scales. Each convolution unit consists of two convolution layers followed by a max pooling and a dropout layer. The outputs from the convolutional units are then concatenated and fed to two fully connected dense layers, each themselves followed by dropout. Finally a softmax activation layer is applied to get a categorical classification.

We trained the `diploS/HIC` CNN using simulations generated as described above. In general both training and independent test sets consisted of 2 *×* 10^3^ simulations per class unless otherwise specified. Optimization of the CNN parameters was performed using the Adam algorithm (Kingma and Ba [2014]) tracking as our objective function the accuracy of the classifier on a validation set. Optimization epochs were run until the validation accuracy stopped improving beyond a difference of 0.001 between epochs. In practice this training was quite rapid, with generally less than 20 epochs needed until stopping.

We implement our CNN using the open source package TensorFlow (Abadi et al. [2016]) and its associated higher level python API Keras (Chollet et al. [2015]). Our software for performing the `diploS/HIC` feature vector calculation and deep learning is available from https://github.com/kern-lab/.

## Results

We were first interested in examining how using a CNN with `diploS/HIC` affected classification accuracy in comparison to the Extra-Trees classifier we had used with `S/HIC`. For this we generated constant population size simulations with moderately strong selection, *α ∼ U* (250, 2500), where the beneficial mutation had fixed immediately prior to sampling (i.e. *τ* = 0) across five different sample sizes of *n* = *{*20, 40, 60, 80, 100*}* haploid chromosomes. The complete parameterization of these simulations is given in Supplemental Table 1. We were also interested in characterizing the difference between using unphased and phased data in both `S/HIC` implementations. Our expectation was that phased data should add significant information to the classification task we are interested in. Figure 5 shows accuracy of four classifiers as a function of sample size. Those classifiers are the original `S/HIC` classifier (labeled “ETC haploid”), a CNN classifier which uses the original `S/HIC` feature vector (“CNN haploid”), and our new, unphased data feature vectors trained with either a CNN (labelled “CNN”) or an ETC (“ETC”). A few trends are notable. First, it is clear that phased data add useful information for classification– both of our phased (haploid) classifiers are more accurate than our our unphased classifiers. Second, our unphased classifiers themselves are performing quite well even at small sample size (for instance *∼* 83% accuracy at *n* = 10 diploid individuals). Third, the convolutional neural network approach adds a bit of accuracy, although the gains aren’t dramatic. While that is so, the size of the training set used here, 2000 examples per class, is quite small for deep learning, thus we might expect to see a greater difference between the CNN and the ETC approaches if we were to increase the size of our training set. These trends generalize to stronger and weaker selection although accuracy varies across these cases (see Figures S2 and S3).

**Figure 5:**
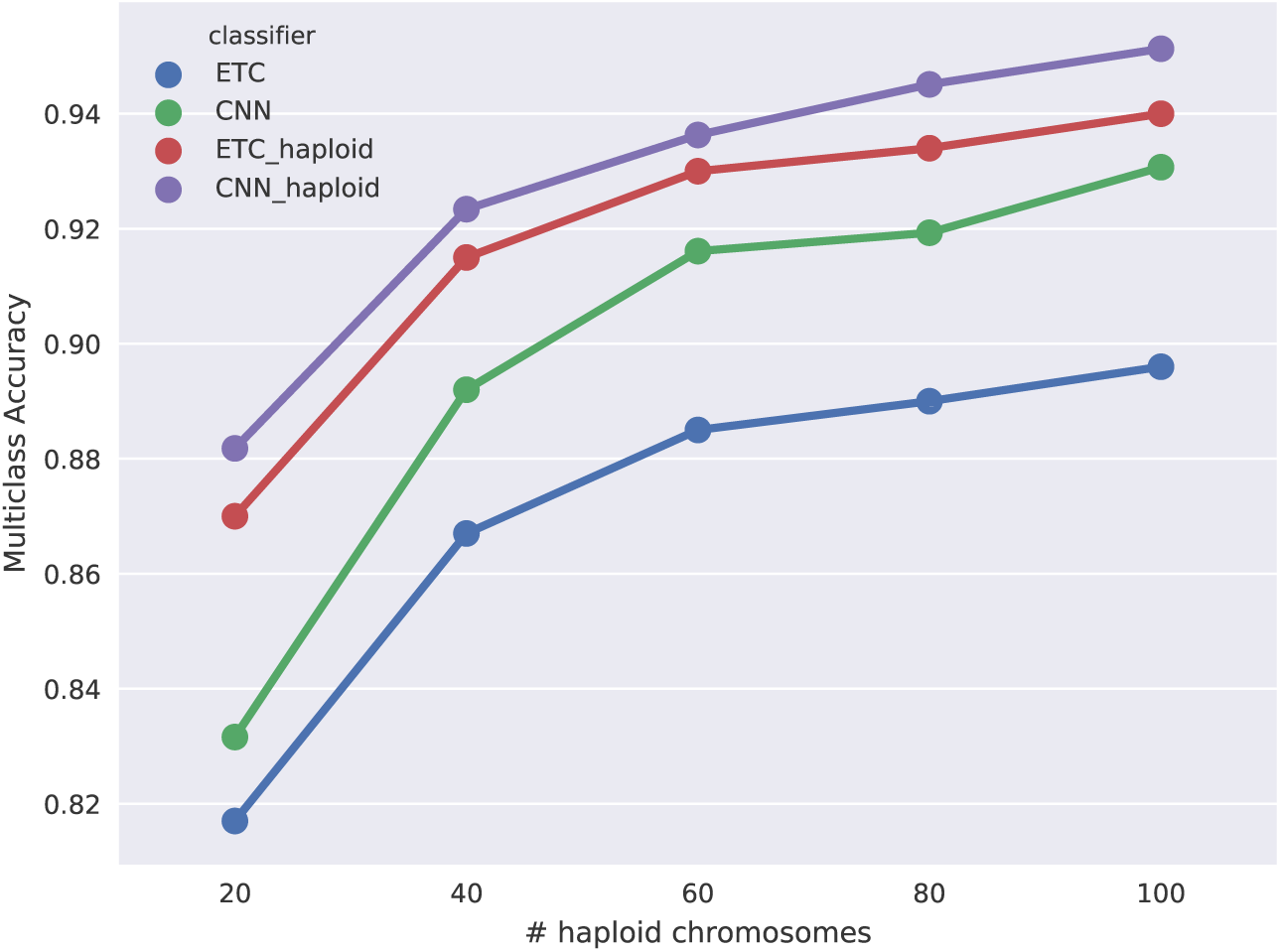
Multiclass classification accuracy for diploS/HIC and related classifiers. Here we compare the multiclass classification accuracy of four related classifiers across a variety of sample sizes: the original `S/HIC` classifier (“ETC haploid”), a `S/HIC` classifier which uses a CNN rather than an ETC (“CNN haploid”), our new classifier for unphased data trained with an ETC (“ETC”), and finally our new classifier training with a CNN (“CNN”). Note the y-axis has been truncated to better display the separation between methods.

We next turn our attention to a slightly more realistic scenario in which the beneficial mutation of interest has not fixed immediately before sampling the population, but instead fixed within some uniform range of time before sampling. We generated simulations as before, under constant population size and moderately strong selection, *α ∼ U* (250, 2500), but allowing the time of fixation, *τ*, to vary such that *τ ∼ U* (0, 0.025). To characterize performance of the classifier under this regime, in Figure 6 we show ROC curves (i.e. true positive vs. false positive rates) for our classifiers’ abilities to distinguish between sweeps, both hard and soft, versus unselected regions, in this case both neutral and linked regions. At all sample sizes we have excellent discriminatory power for this binary classification task. This is also the case for stronger and weaker selection (Figures S4 and S5). Across each of these scenarios considered `diploS/HIC` is surprisingly competitive with our original, phased data implementation. This suggests that for sweep finding `diploS/HIC` will be a very useful tool in datasets without information on phase. ROC curves show one aspect of our `diploS/HIC` classifier, but perhaps more illuminating is the examination of classification in a spatial context (i.e. along a recombining chromosome). To do this we visualize the confusion matrix for 10 subwindows surrounding a central subwindow that has undergone either a hard or soft sweep. Simulated subwindow examples are then classified as belonging to one of five classes (hard sweep, soft sweep, linked to a hard sweep, linked to a soft sweep, or neutral) and the fraction of each such classification is recorded. Figure 7 shows such a confusion matrix for a constant size population with moderately strong selective sweeps, *α ∼ U* (250, 2500), and for an intermediate sample size of *n* = 60 haploid chromosomes. For this sample size and parameterization `diploS/HIC` performs quite similarly to our original haploid `S/HIC` implementation with greater than 86% of sweep windows identified correctly and with a false positive rate of *∼* 5%. The original `S/HIC` had a both a slightly higher accuracy on sweeps (*∼* 87.5%) and false positive rate (*∼* 6.7%) on the same test scenario (but with a slightly smaller sample size of *n* = 50; see Schrider and Kern [2016] Figure 4).

**Figure 6:**
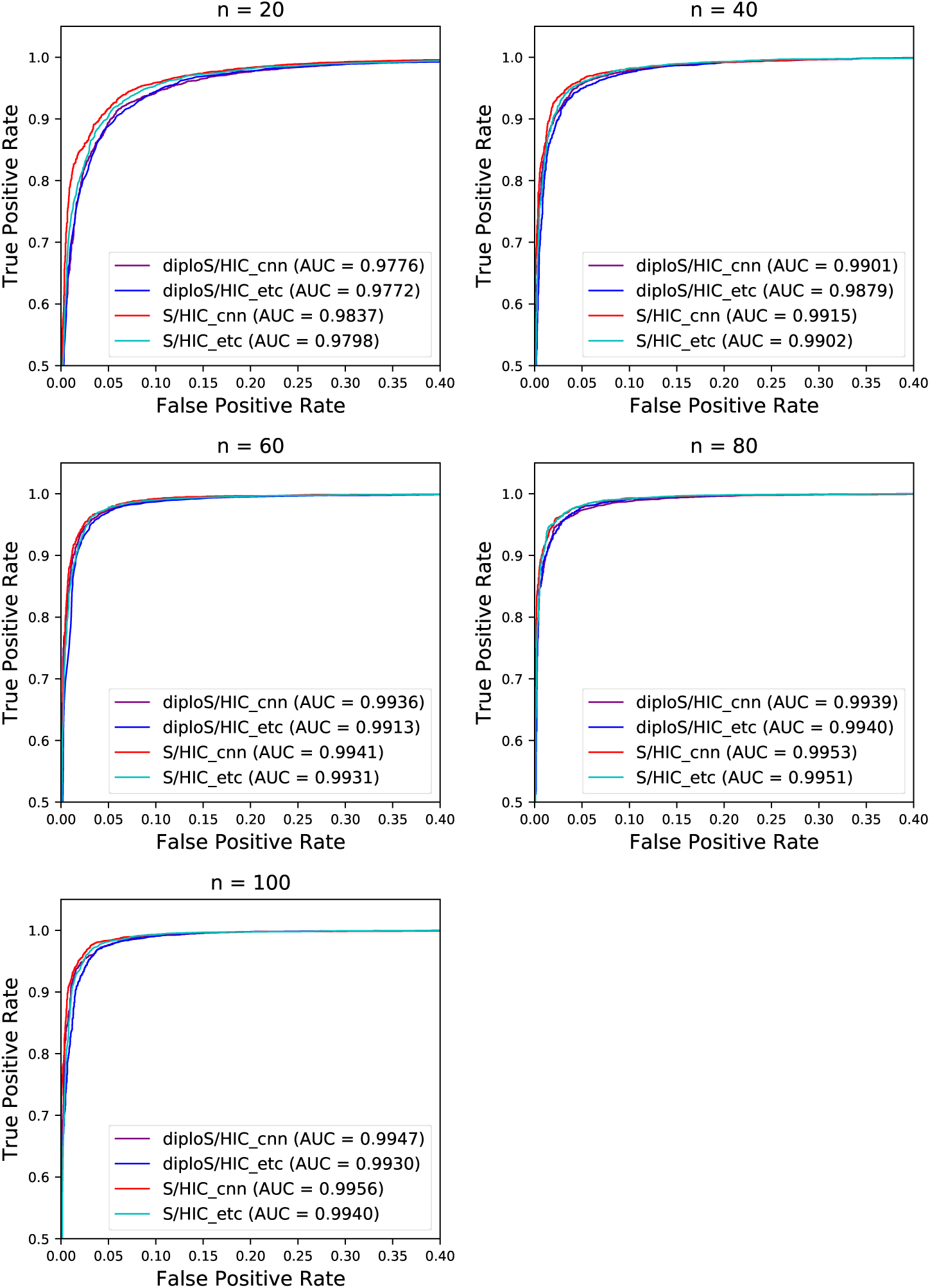
ROC curves showing the true and false positive rates for various classifiers. Here we show ROC curves for the binary classification task of Hard and Soft vs linked and neutral regions. ROC curves are shown for both unphased and phased data classifiers as well as for the two machine learning algorithms considered. Each panel represents a different sample size of haploid chromosomes from *n* = 20, 40, 60, 80, 100. Note the scale of the x and y axes does not run from (0,1) but instead is zoomed in to show what little separation among methods there is.

**Figure 7:**
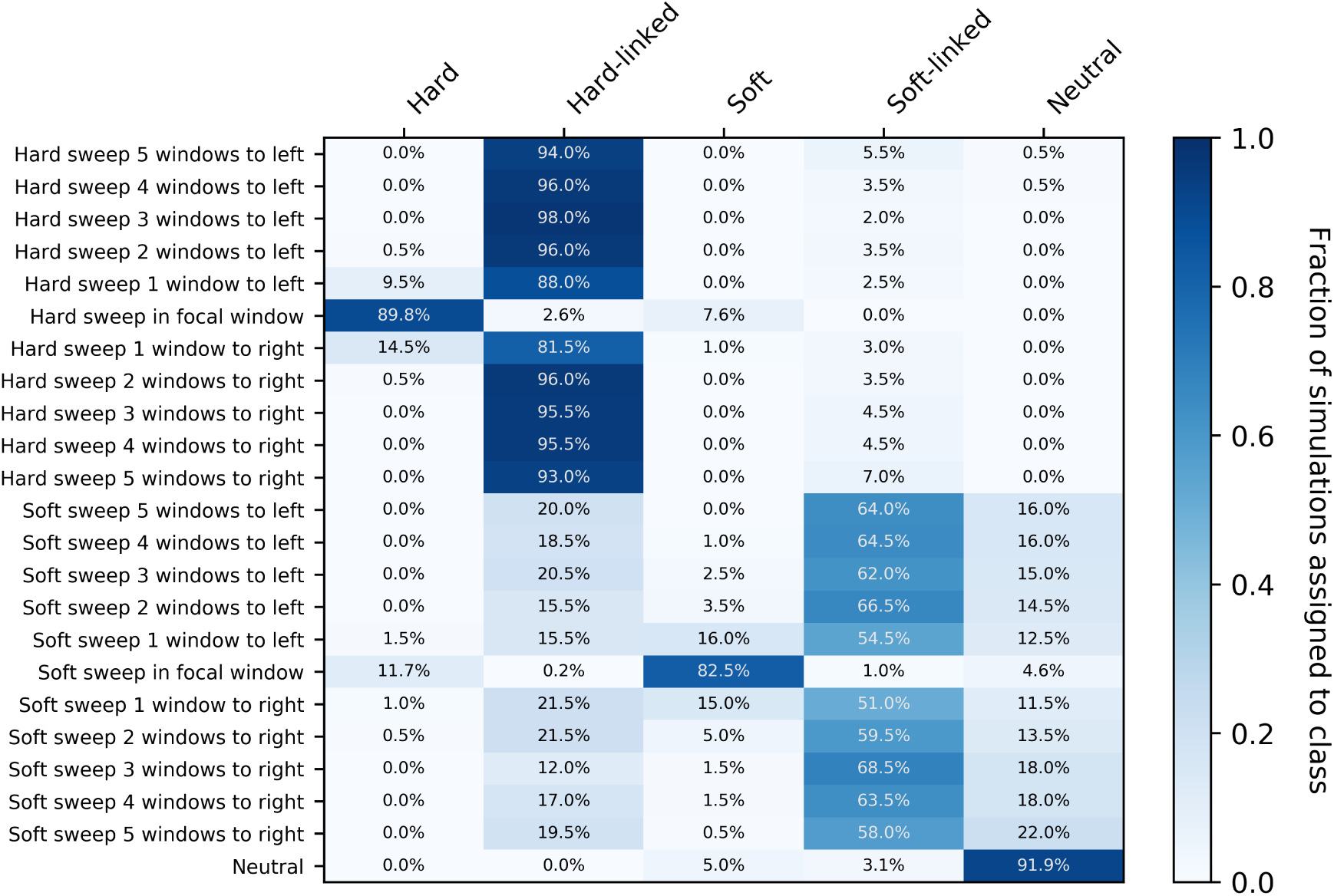
Confusion matrices of subwindows across a recombining chromosome. On the y-axis the location of the classified subwindow relative to the sweep is shown while the x-axis shows the predicted class for each subwindow from `diploS/HIC`. These results are from simulations using the same parameter set as in Figure 6, using a sample size of *n* = 60 haploid chromosomes

For stronger sweeps we see qualitatively similar patterns, with an excellent ability to localize both hard and soft selective sweeps (Figure 8). Simulating sweeps with *α ∼ U* (2500, 25000) but holding constant the rate of recombination means that a greater portion of the chromosome is affected by the sweep. We can see this in our increased misclassification rate of subwindows closest to hard sweeps in Figure 8 relative to Figure 7. Nevertheless our multiclass accuracy (the prediction accuracy across all data classes) on stronger sweeps is still high (*∼* 94%; Figure S2).

**Figure 8:**
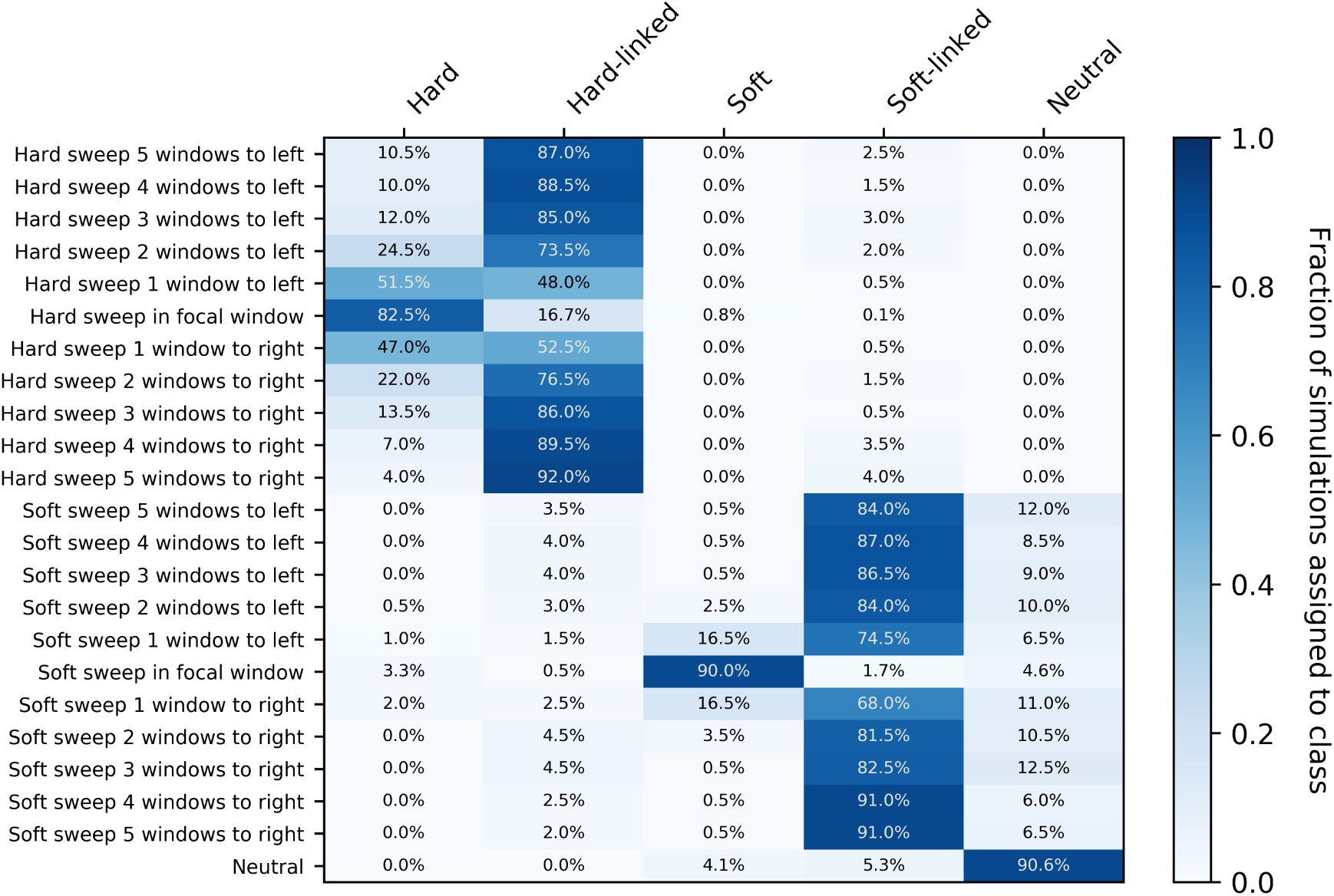
Confusion matrices of subwindows across a recombining chromosome with stronger selection. On the y-axis the location of the classified subwindow relative to the sweep is shown while the x-axis shows the predicted class for each subwindow from `diploS/HIC`. These results are from simulations using the same parameter set as in Figure 6 with the exception of selection which is an order of magnitude stronger, *α U* (2500, 25000), and again using a sample size of *n* = 60 haploid chromosomes

On the other hand, when selection is comparatively weak, a relatively smaller portion of the chromosome is perturbed by a selective sweep and thus sweep signatures can be harder to detect, but somewhat easier to localize if detected. In Figure S6 we show confusion matrices associated with an order of magnitude weaker selection, using *α ∼ U* (25, 250). In this scenario we have reduced sensitivity to sweeps, a slightly higher false positive rate, and struggle to determine the mode of selection. With weak selection `diploS/HIC` is less likely to identify flanking regions as sweeps, though we struggle with classification of hard-linked and soft-linked regions, misclassifying each type both as the other, or often as neutral in the case of soft-linked sites; such errors are of minor concern if one’s primary goal is the discrimination between sweeps and unselected regions. Supporting this point is the fact that our binary classifier (sweep vs. linked+neutral) has a high area under the curve (AUC; *∼* 0.96) even though our multiclass accuracy is lower (*∼* 64%; see Figure S3); thus even with very weak selection our classifier can find sweeps.

Each of the above scenarios simulated populations with constant population size, which is a rarity in the natural world. We thus turn attention to the more realistic case in which population size has changed in recent history. In particular we sought to characterize the performance of `diploS/HIC` for samples from populations with strong growth, such as we recently observed in a Burkino Faso population of *Anopheles gambiae* (Anopheles gambiae 1000 Genomes Consortium et al. [2017]). We inferred using information from the site frequency spectrum that this sample, shorthanded BFS, had experienced greater than 30-fold growth in the past 100,000 years (Anopheles gambiae 1000 Genomes Consortium et al. [2017]). We simulated training sets under the BFS sample’s inferred demographic parameters (see Supplemental Table 1) for a variety of sample sizes and selection scenarios. Figure 9 shows the multiclass accuracy of `diploS/HIC` (labeled CNN) and related classifiers when testing against an independent set of simulations.

**Figure 9:**
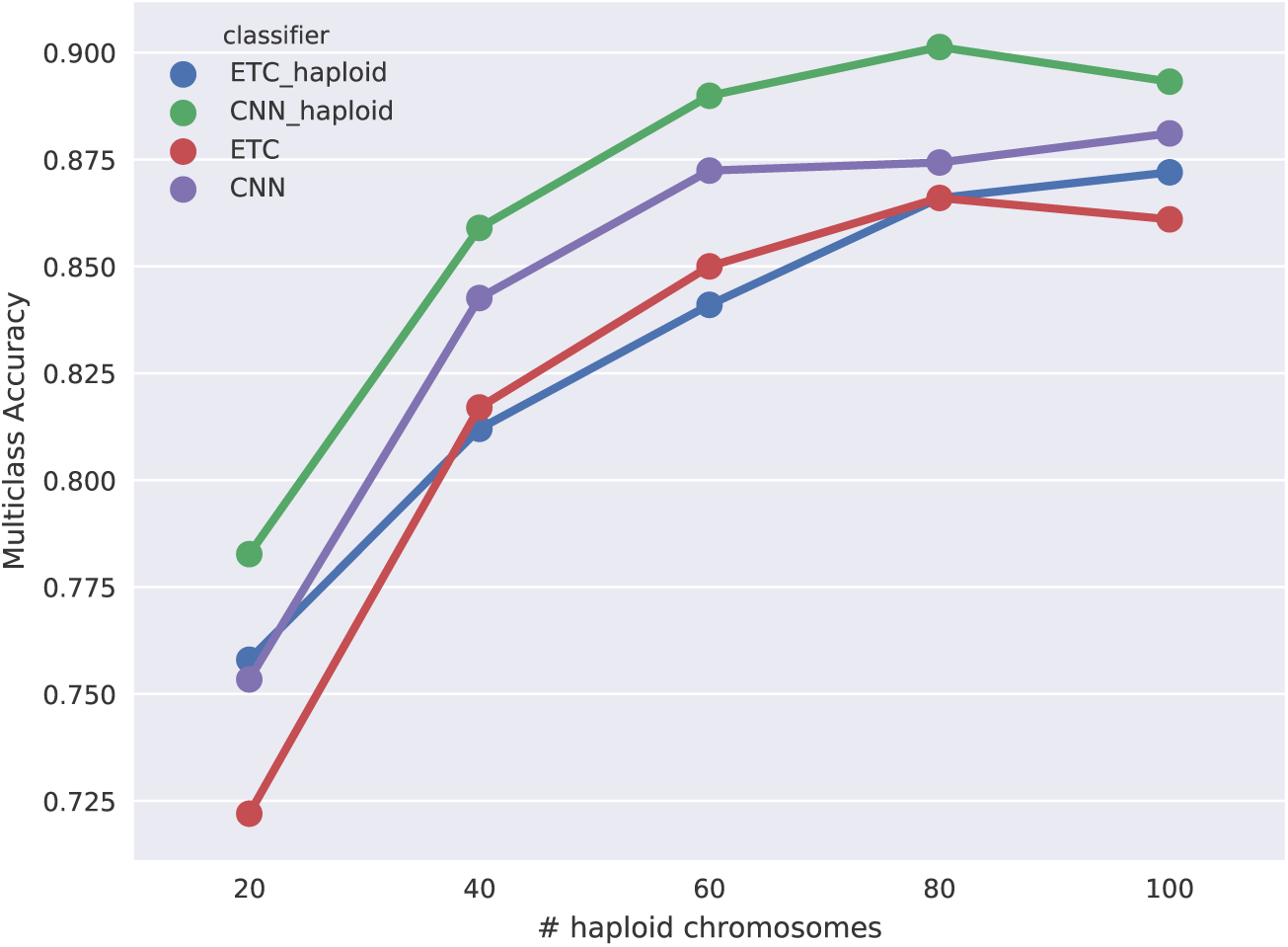
Multiclass classification accuracy for diploS/HIC and related classifiers for *Anopheles* demographic history with moderate selection. Here we compare the multiclass classification accuracy of four related classifiers across a variety of sample sizes: the original `S/HIC` classifier (“ETC haploid”), a `S/HIC` classifier which uses a CNN rather than an ETC (“CNN haploid”), our new classifier for unphased data trained with an ETC (“ETC”), and finally our new classifier training with a CNN (“CNN”).

As in the case of constant population size, the general trend is that CNN-based classifiers consistently outperform those based on Extra-Trees. Moreover in this case `diploS/HIC` even tends to outperform our original implementation of `S/HIC` which uses phased haplo-types. Supplementary Figures S7 and S8 show the same comparisons for when selection is an order of magnitude stronger and weaker respectively. Under this demographic scenario in our stronger selection case, *α ∼ U* (25000, 250000), selection is strong relative to recombination and we lose the ability to localize hard sweeps, resulting in a loss of accuracy. On the contrary our weak selection scenario for *Anopheles* yields excellent multiclass accuracies. As before, we prefer to visualize performance using confusion matrices, and in Figure 10 we show representative classifications as a function of distance from each sweep type for simulations of BFS demography and moderately strong sweeps. With *n* = 60 haploid chromosomes we have 87% accuracy and our ability to localize sweeps is quite good, although selection in this case is sufficently strong relative to recombination such that we have a significant misclassification rate of windows neighboring hard sweeps into the hard sweep category. If selection is an order of magnitude weaker for BFS demography then our localization for those windows neighboring sweeps improves considerably (Figure S10). However, if selection is an order of magnitude stronger `diploS/HIC` and related classifiers all struggle to accurately find hard sweeps, as for this parameterization selection is too strong relative to recombination (Figure S9), therefore larger window sizes would be required.

**Figure 10:**
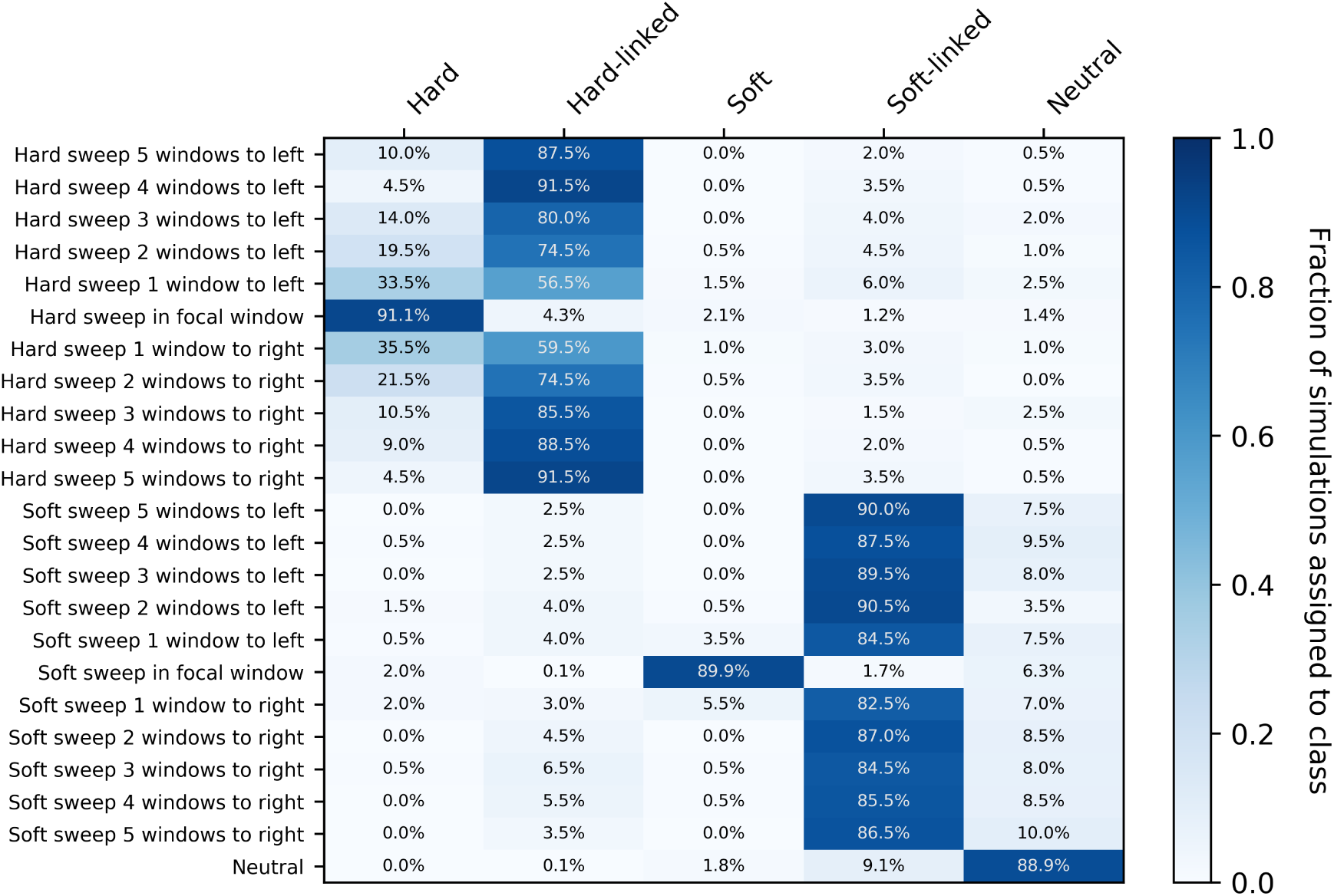
Confusion matrices of diploS/HIC for an *Anopheles gabiae* demographic history. On the y-axis the location of the classified subwindow relative to the sweep is shown while the x-axis shows the predicted class for each subwindow from `diploS/HIC`. These results are from simulations using the parameters reflecting the population history of a BFS sample of *Anopheles gambiae*, using a sample size of *n* = 60 haploid chromosomes. Here we have simulated sweeps with moderate selection coefficients, *α U* (250, 2500)

We previously showed our `S/HIC` classifier to be quite robust to demographic model misspecification (Schrider et al. [2016]), thus we were curious to see if our new implementation should share similar properties. *A priori* we expect `diploS/HIC` to be robust, as such robustness stems from the summary statistic transform that we apply to our input statistics, rather than some higher level property of the ML algorithm. To characterize accuracy in the face of model misspecification we used the classifiers that we had trained on constant size population simulations (e.g. Figure 5) to make predictions from data drawn from the BFS sample demographic model (i.e. strong population growth). In Figure S11 we show ROC curves for both our original `S/HIC` and `diploS/HIC` with and without model misspecification for each of a set of five sample sizes. Both `S/HIC` and `diploS/HIC` are indeed robust to model misspecification and while `S/HIC` is more accurate, owing to its use of haploid information, `diploS/HIC` is more robust to such misspecification.

Practicing population geneticists often are faced with data can only be phased computationally, rather than through transmission or inbreeding. We were thus interested in characterizing the performance of our classifiers in the face of phasing switch errors. To explore this, we trained haploid and diploid `diploS/HIC` classifiers, along with `S/HIC` classifiers on data that contained no switch errors, but then tested those trained classifiers on data which had switch errors inserted at various probabilities (i.e. genotypes at heterozygous sites were flipped at a given rate). In Figure S12 we show the accuracy of three classifiers as a function of switch error rate. Those classifiers are the original `S/HIC` classifier (labeled “ETC haploid”), a CNN classifier which uses the original `S/HIC` feature vector (“CNN haploid”), and our unphased `diploS/HIC` classifier (labelled “CNN”). As expected the accuracy of the unphased classifier does not change with increasing switch error rate, however both of the classifiers that make use of phase information do lose accuracy. Thus if there is reason to believe that computational phasing might lead to appreciable error rates we recommend that the user should use the default, unphased version of `diploS/HIC`.

In general we find that under a more complex history of population growth, such as that from the BFS sample, `diploS/HIC` performs quite well at finding both hard and soft sweeps and differentiating among them. Moreover we find that `diploS/HIC` is quite robust to model misspecification during training. This is the case even for reasonably small population sizes, although our smallest sample size tested did lag in accuracy considerably. Thus a fair degree of caution should be taken when analyzing smaller samples (i.e. *n ≤* 20) using our method.

### An application to *Anopheles gabiae* genome sequence data

As an example of how `diploS/HIC` might be used to elucidate the evolutionary history of an organism we here provide a targeted look at the *Gste* region of the *A. gambiae* genome. Over the last 20 years as part of the Roll Back Malaria initiative, mosquito populations have been subject to large-scale application of insecticides and as a consequence have begun to adapt to this new environment through the evolution of resistance (Hemingway et al. [2016]). The *Gste* locus contains a cluster of glutathione S-transferase genes including *Gste2*, which has previously been shown to be involved in detoxification of pyrethroids and DDT (Mitchell et al. [2014]). In Figure 11 we show `diploS/HIC` classifications of windows throughout the region at three physical scales (10kb, 20kb, and 50kb). For computational convenience, we trained a classifier using simulations of genetic distances that correspond to *∼*5kb, but predict on feature vectors from a larger region. Though ideally simulation with a matching window size should be used, this is acceptable for our purposes here because our feature vector itself is scale free, and the ratio of the recombination rate to the selection coefficient is what determines the spatial patterns of variation across a region. At each scale a soft sweep is identified at the *Gste* cluster, and one can see from patterns of Tajimas’s *D*, *J*_12_, and *ZnS*, that the signal of selection extends for a long distance in each direction away from the sweep. This squares nicely with the haplotype based analysis done by Anopheles gambiae 1000 Genomes Consortium et al. [2017] which showed that numerous haplotypes were under selection at *Gste*. We have included the workflow of this entire empirical application as an example use case of `diploS/HIC` here: https://github.com/kern-lab/diploSHIC/wiki/A-soup-to-nuts-example

**Figure 11:**
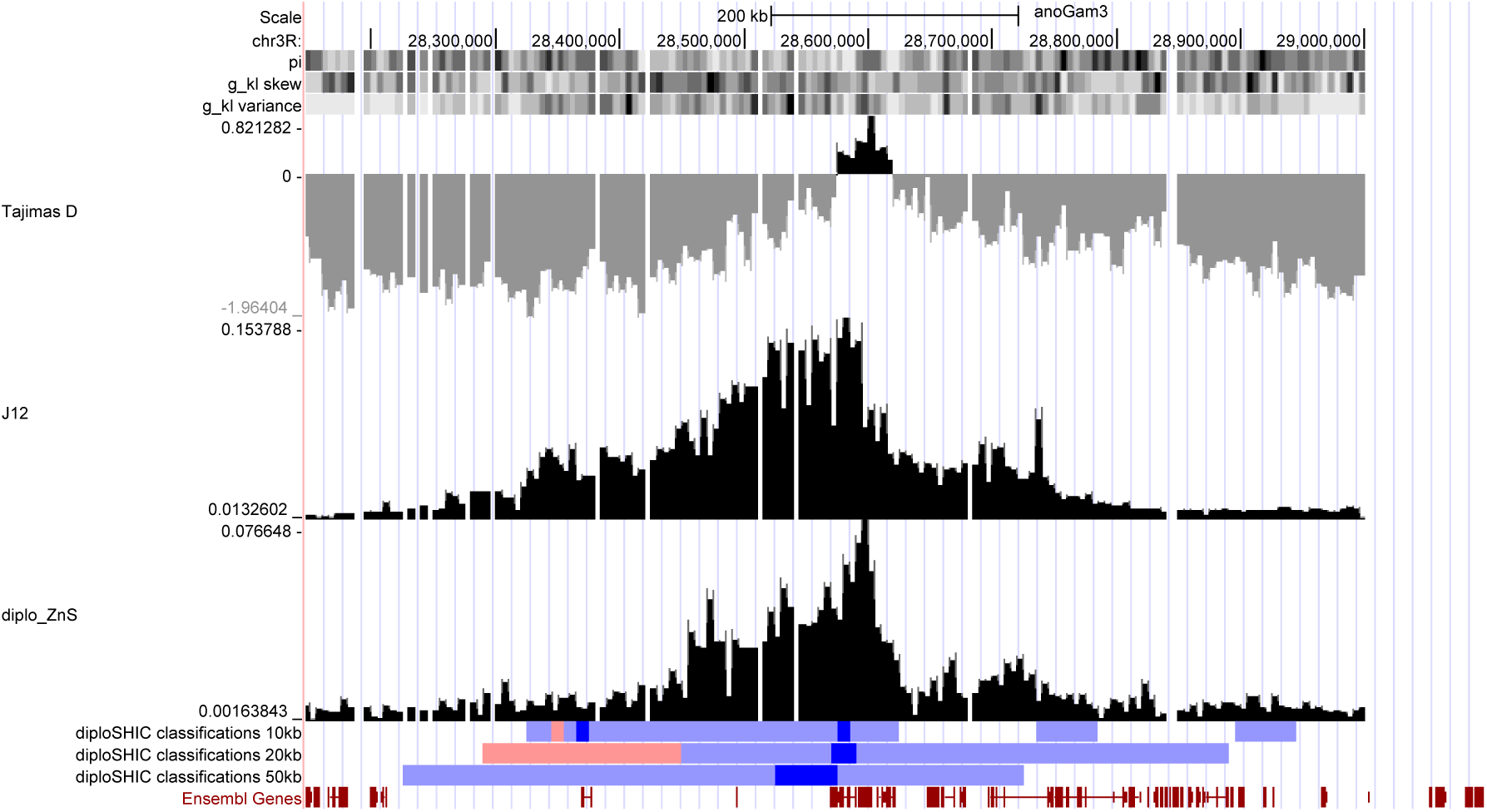
A large soft sweep at *Gste* in *Anopheles gambiae*. Here we show `diploS/HIC` classifications at three physical scales along with associated summary statistics for a megabase region surrounding the *Gste* cluster on chromosome 3R for the BFS sample. `diploS/HIC` classifications are colored here such that windows classified as soft sweeps are dark blue, linkedSoft as light blue, and linkedHard as light red. Above the `diploS/HIC` classifications we show a subset of the summary statistics used as part of the feature vector (see Methods for details).

## Discussion

Creating a complete catalog of selective sweeps in the genome of any organism remains an elusive goal. Population genetics, as a field, requires knowledge of the targets of selection for at least two separate purposes. At a broad scale, we would like to know about genome-wide parameters such as the rate of selective sweeps per generation and their associated distribution of selective effects. At a finer scale, we would like to know the precise location of individual beneficial fixations, so that we can learn about the functional ramifications of adaptive evolution. Thus through the creation of a catalog of selective sweeps we hope to learn both about the process of adaptation as well as the nature of the adaptations themselves.

While methods development for sweep finding tools has a rich history in the literature, in the past few years substantial performance gains have come through leveraging supervised machine learning approaches (reviewed in Schrider and Kern [2018]). In particular, the use of machine learning has led to more powerful methods for detecting sweeps that are robust to complicated demographic histories which had confounded earlier approaches. While this is so, nearly all machine learning methods for finding selection require the use of phased haplotype information for input (the sole exception being SF-Sselect (Ronen et al. [2013]), which is based strictly on the site frequency spectrum). In this report we present a new method that we call `diploS/HIC` that has exceptional power for finding both hard and soft selective sweeps and differentiating among them without requiring phased data. `diploS/HIC` builds on the strategy of our earlier method, `S/HIC` (Schrider and Kern [2016]), in utilizing a large vector of summary statistics that have been transformed to capture spatial information around a focal region to be classified. For `diploS/HIC` we have introduced a number of new population genetic summary statistics that are compatible with unphased genotypes and which are useful for detecting and discriminating between hard and soft selective sweeps. Note that, like the original `S/HIC`, `diploS/HIC` was designed with completed sweeps in mind, but could be extended to handle incomplete sweeps in a straightforward manner by simulating appropriate training data and including additional summary statistics.

Algorithmically `diploS/HIC` utilizes a deep convolutional neural network (CNN) as the basis of its classification (LeCun et al. [2004], Krizhevsky and Hinton [2010], Krizhevsky et al. [2012]). Thus, we cast the sweep finding problem as one of image recognition. Our goal in doing so is to more explicitly utilize the spatial covariance structure of summary statistics in a given region, beyond the simple spatial transformation we had been using in Schrider and Kern [2016]. While it seems this has succeeded to some degree (i.e. we see that a CNN classifier using the `S/HIC` feature vector outperforms our original implementation – Figure 5) we suspect that there is still considerable room for improvement. Firstly, the use of recurrent neural network approaches that are specifically formulated for capturing signals in sequential data should provide a principled way of gleaning further spatial signal using summary statistics (Graves et al. [2013], Sutskever et al. [2014]). Secondly, we imagine that improvements in using the full information in the spatial arrangement of polymorphism surrounding a sweep could come from abandoning summary statistic representations of loci, and instead using image recognition (i.e. CNN) approaches on images created directly from sequence alignments of a genomic region (Chan et al. [2018]; L. Flagel, *pers. comm.*). A similar approach using images of read pileups was recently introduced for variant calling, yielding solid performance improvements over competing methods (Poplin et al. [2017]).

In summary we have shown that `diploS/HIC` performs quite well when compared to our previous `S/HIC` under both simple and more complex population size histories. Identifying selective sweeps in populations with non-equilibrium demographic histories remains an important and difficult problem, particularly in cases where the underlying demographic model is unknown or poorly estimated. For instance it is well known that certain demographic models, such as population growth, can mimic selective perturbations (Simonsen et al. [1995], Jensen et al. [2005]). Here we have shown as with `S/HIC` before it, `diploS/HIC` is accurate in the face of non-equilibrium demography, even when misspecified. In conclusion, we believe that `diploS/HIC` provides yet another powerful tool for population geneticists to use when phased information is unavailable.

## Acknowledgements

We thank Jeff Adrion, Jeff Ross-Ibarra, and Alexander Xue for comments on this manuscript. ADK was supported by National Institute of Health award no. R01GM117241. DRS was supported by NIH award no. K99HG008696.

## BFS demography string

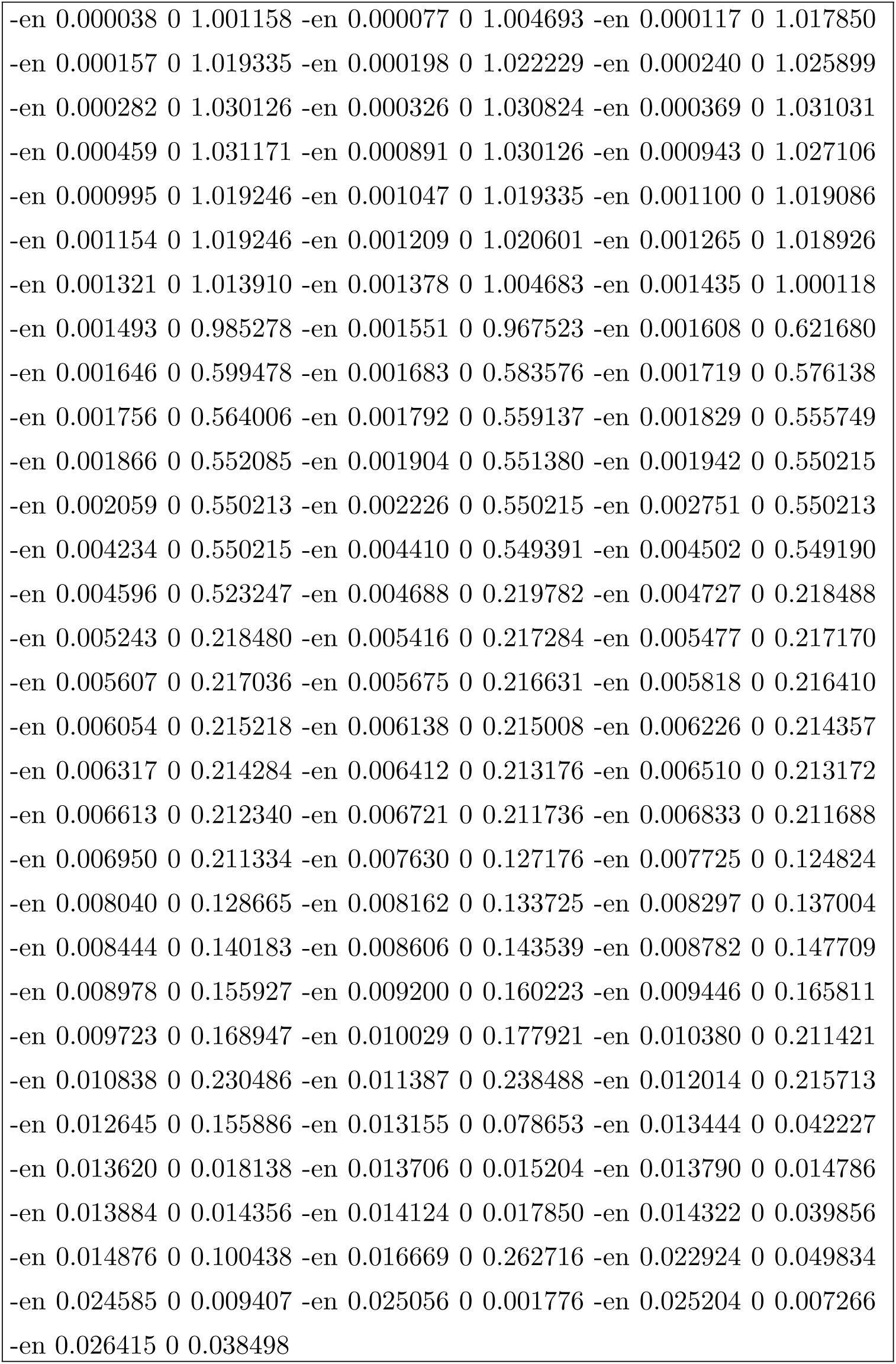

**Table S1:** Parameters used for simulating training and test datasets

|  | Equilibrium<br>(weak selection) | Equilibrium<br>(moderate selection) | Equilibrium<br>(strong selection) | Burkino Faso Demography<br>(moderate selection) | Burkino Faso Demography<br>(strong selection) | Burkino Faso Demography<br>(weak selection) |
| --- | --- | --- | --- | --- | --- | --- |
| Mutation rate<br>(4NL, where L is the<br>length of the simu-<br>lated chromosome) | $\sim U(528, 1584)$ | $\sim U(528, 1584)$ | $\sim U(528, 1584)$ | $\sim U(1750.2, 17502.0)$ | $\sim U(1750.2, 17502.0)$ | $\sim U(1750.2, 17502.0)$ |
| Recombination rate<br>(4NrL) | 880 | 880 | 880 | $\sim U(19252.2, 57756.8)$ | $\sim U(19252.2, 57756.8)$ | $\sim U(19252.2, 57756.8)$ |
| Selection coefficient<br>for sweeps (2Ns) | $\sim U(25, 2.5 \times 10^2)$ | $\sim U(2.5 \times 10^2, 2.5 \times 10^3)$ | $\sim U(2.5 \times 10^3, 2.5 \times 10^4)$ | $\sim U(2.5 \times 10^3, 2.5 \times 10^4)$ | $\sim U(2.5 \times 10^4, 2.5 \times 10^5)$ | $\sim U(2.5 \times 10^2, 2.5 \times 10^3)$ |
| Fixation time of sweep | $\sim U(0, 0.05)$ | $\sim U(0, 0.05)$ | $\sim U(0, 0.05)$ | $\sim U(0, 4 \times 10^{-5})$ | $\sim U(0, 4 \times 10^{-5})$ | $\sim U(0, 4 \times 10^{-5})$ |
| Initial selected fre-<br>quency of soft sweep | $\sim U(0.0, 0.2)$ | $\sim U(0.0, 0.2)$ | $\sim U(0.0, 0.2)$ | $\sim U(0.0, 0.2)$ | $\sim U(0.0, 0.2)$ | $\sim U(0.0, 0.2)$ |
| Example command<br>line for neutral simu-<br>lation | discoal 100 200 110000<br>-Pt 528 1584 -r 880 | discoal 100 200 110000 -<br>Pt 528 1584 -r 880 | discoal 100 200 110000 -<br>Pt 528 1584 -r 880 | discoal 20 200 55000 -Pt<br>1750.204699 17502.046985 -<br>Pre 19252.251684 57756.755051<br><BFS_Demography>* | discoal 20 200 55000 -Pt<br>1750.204699 17502.046985 -<br>Pre 19252.251684 57756.755051<br><BFS_Demography>* | discoal 20 200 55000 -Pt<br>1750.204699 17502.046985 -<br>Pre 19252.251684 57756.755051<br><BFS_Demography>* |

BFS\_Demography is given in a the preceding supplement subsection

**Figure S1:**
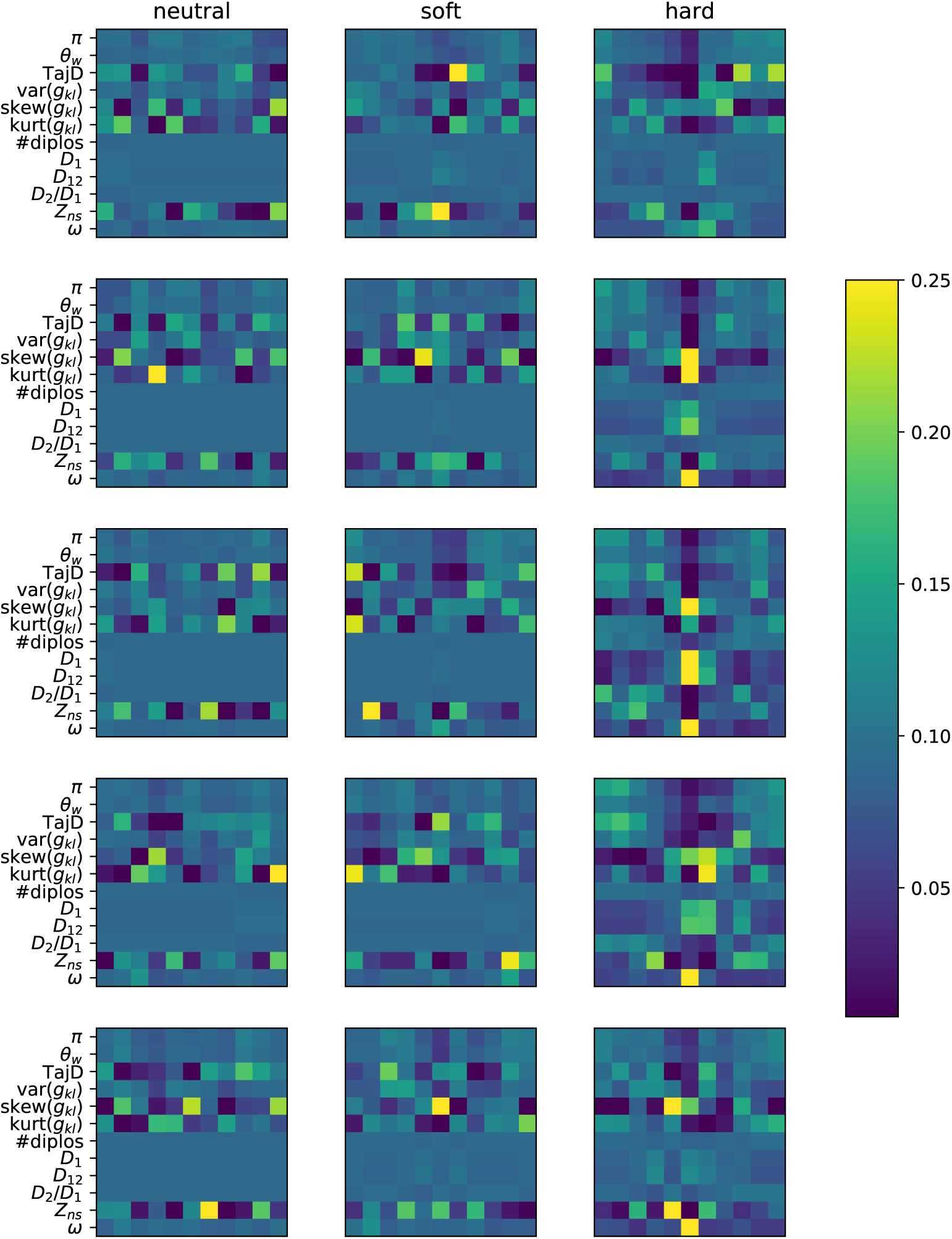
Image instances from single simulations of three classes. Here we should representative images from individual simulations of neutral loci, soft sweeps, and hard sweeps in the left, center, and right columns respectively. Each image represents a different simulation. These simulations were performed under constant population size with parameters according to Figure 1.

**Figure S2:**
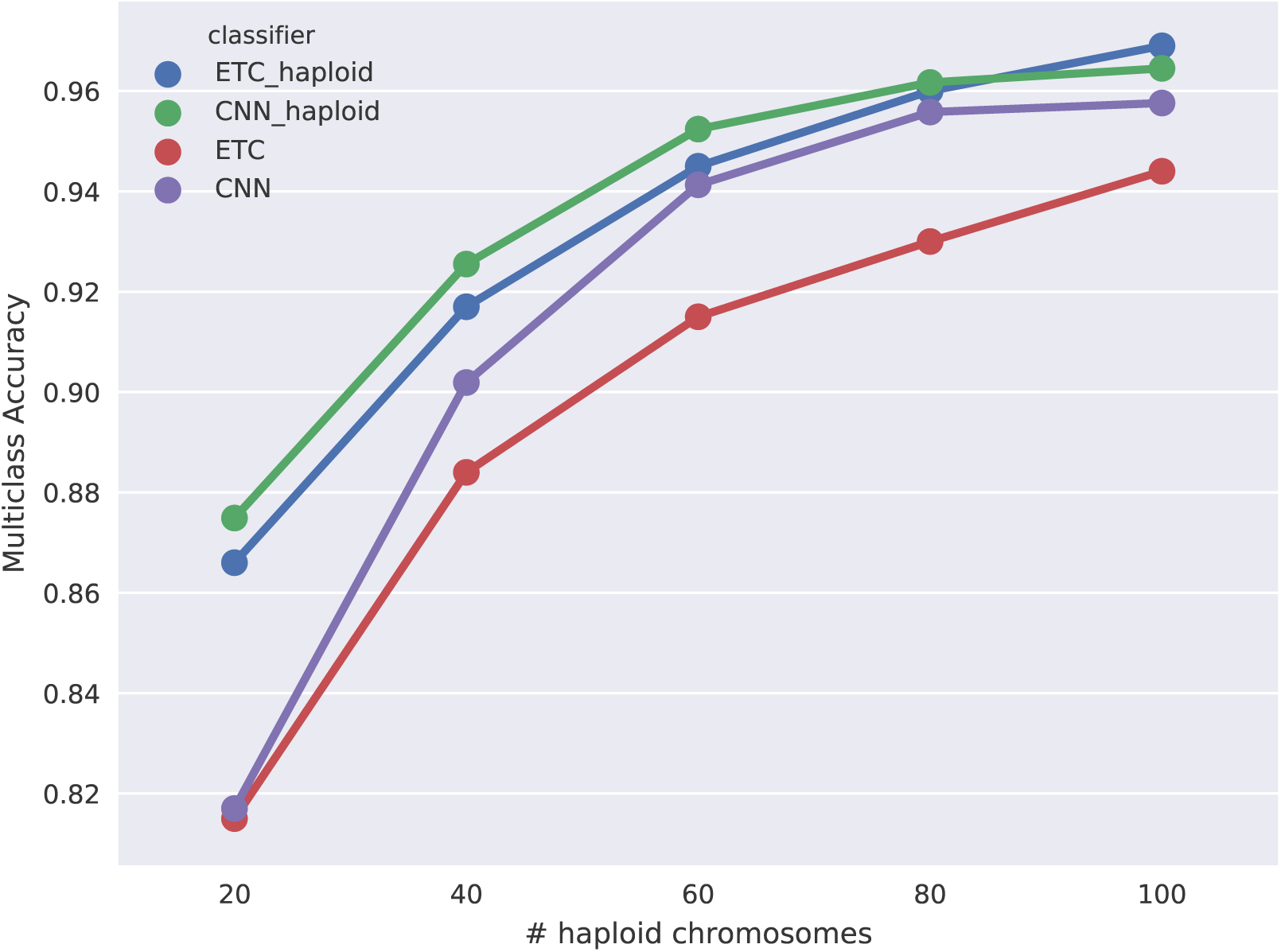
Multiclass classification accuracy for diploS/HIC and related classifiers with strong selection. Here we compare the multiclass classification accuracy of four related classifiers: the original `S/HIC` classifier (“ETC haploid”), a `S/HIC` classifier which uses a CNN rather than an ETC (“CNN haploid”), our new classifier for unphased data trained with an ETC (“ETC”), and finally our new classifier training with a CNN (“CNN”). Simulation parameters are as in Figure 5 but with stronger selection, *α* ~ *U*(2500; 25000)

**Figure S3:**
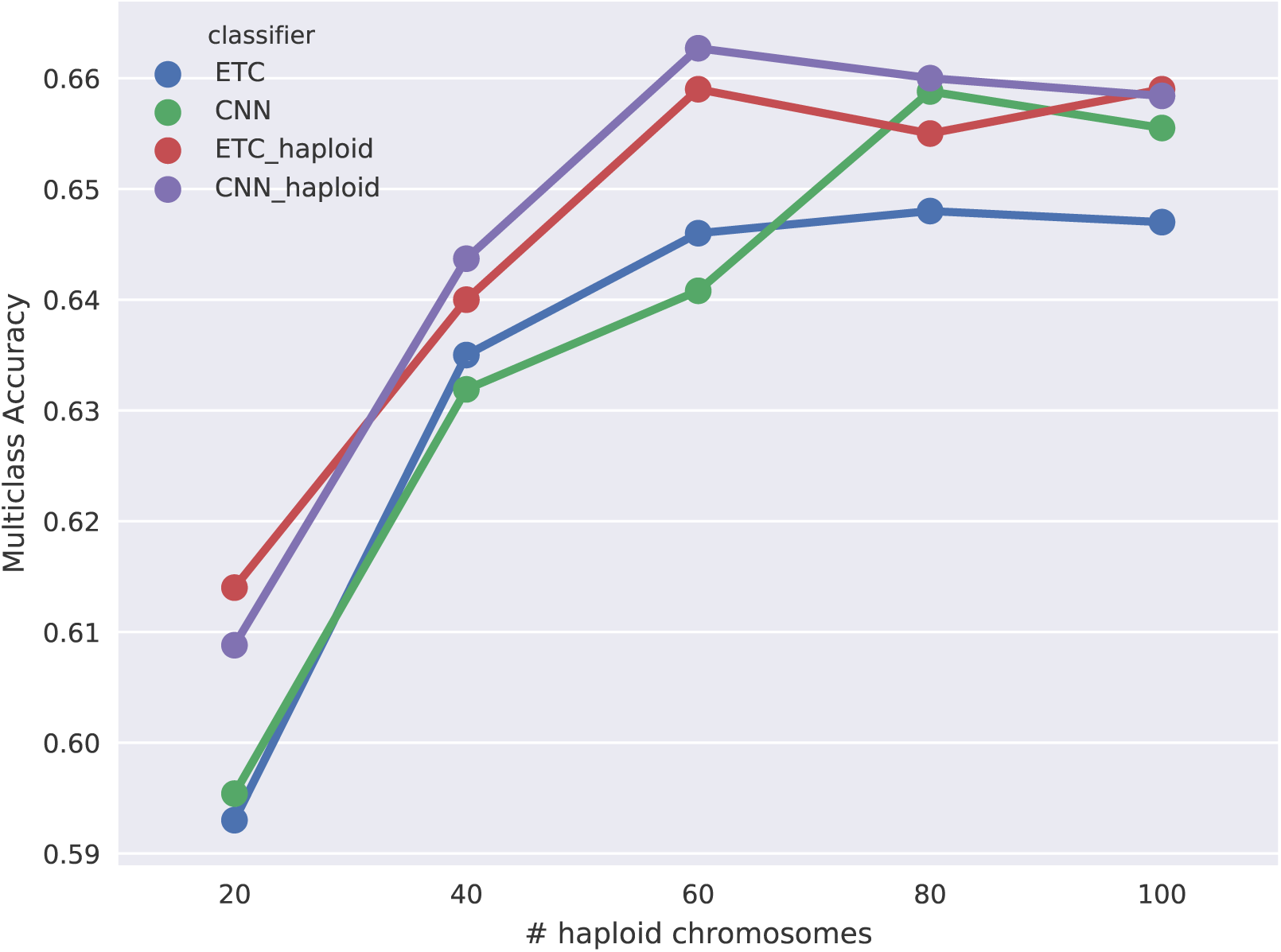
Multiclass classification accuracy for diploS/HIC and related classifiers with weak selection. Here we compare the multiclass classification accuracy of four related classifiers: the original `S/HIC` classifier (“ETC haploid”), a `S/HIC` classifier which uses a CNN rather than an ETC (“CNN haploid”), our new classifier for unphased data trained with an ETC (“ETC”), and finally our new classifier training with a CNN (“CNN”). Simulation parameters are as in Figure 5 but with stronger selection, *α ∼ U* (25, 250)

**Figure S4:**
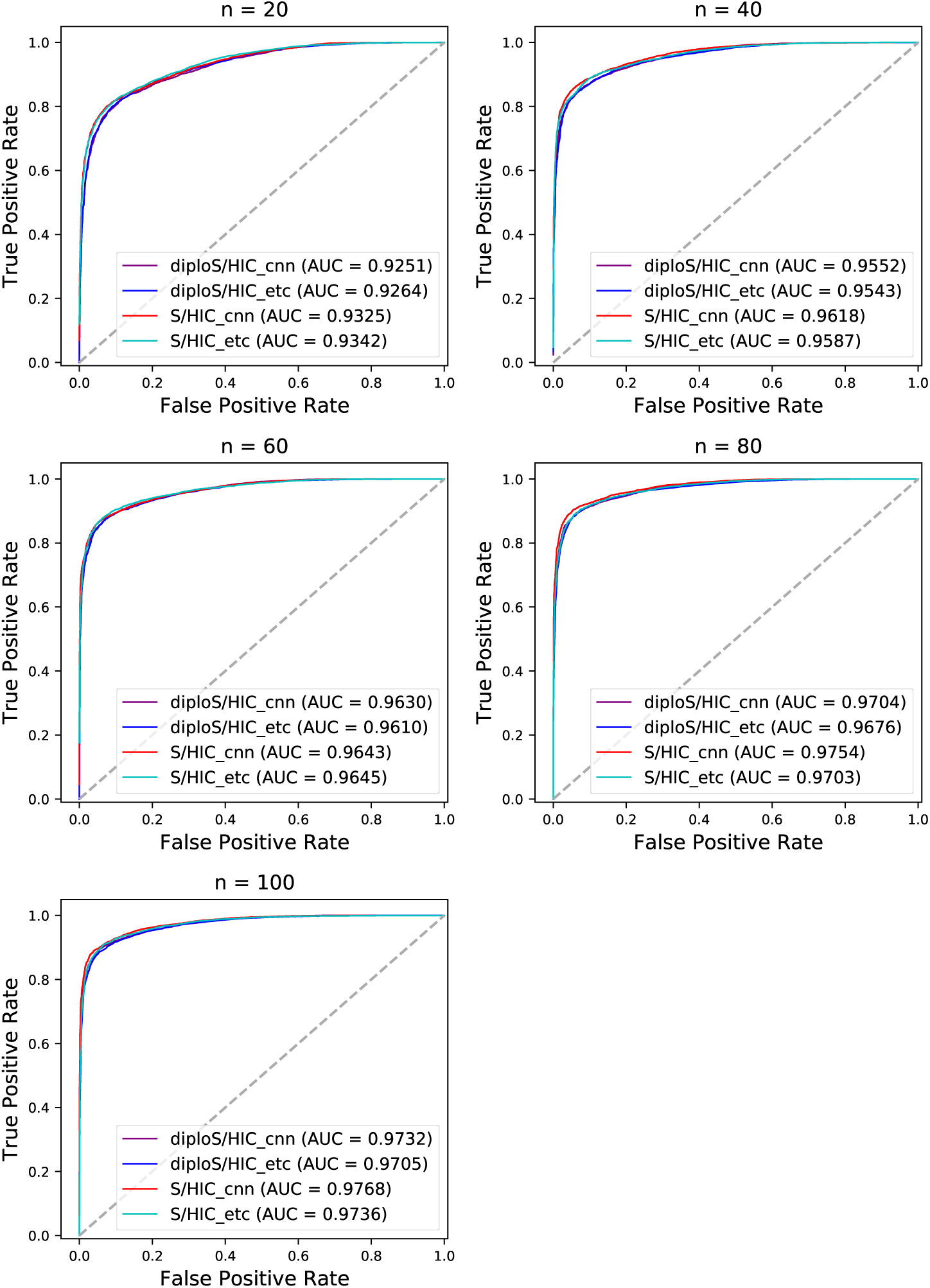
ROC curves showing the true and false positive rates for various classifiers with strong selection. Here we show ROC curves for the binary classification task of Hard and Soft vs linked and neutral regions. ROC curves are shown for both unphased and phased data classifiers as well as for the two machine learning algorithms considered. These simulations were run with the same parameters as those in Figure 6 but with stronger selection, *α ∼ U* (2500, 25000) Each panel represents a different sample size of haploid chromosomes from *n* = *{*20, 40, 60, 80, 100*}*.

**Figure S5:**
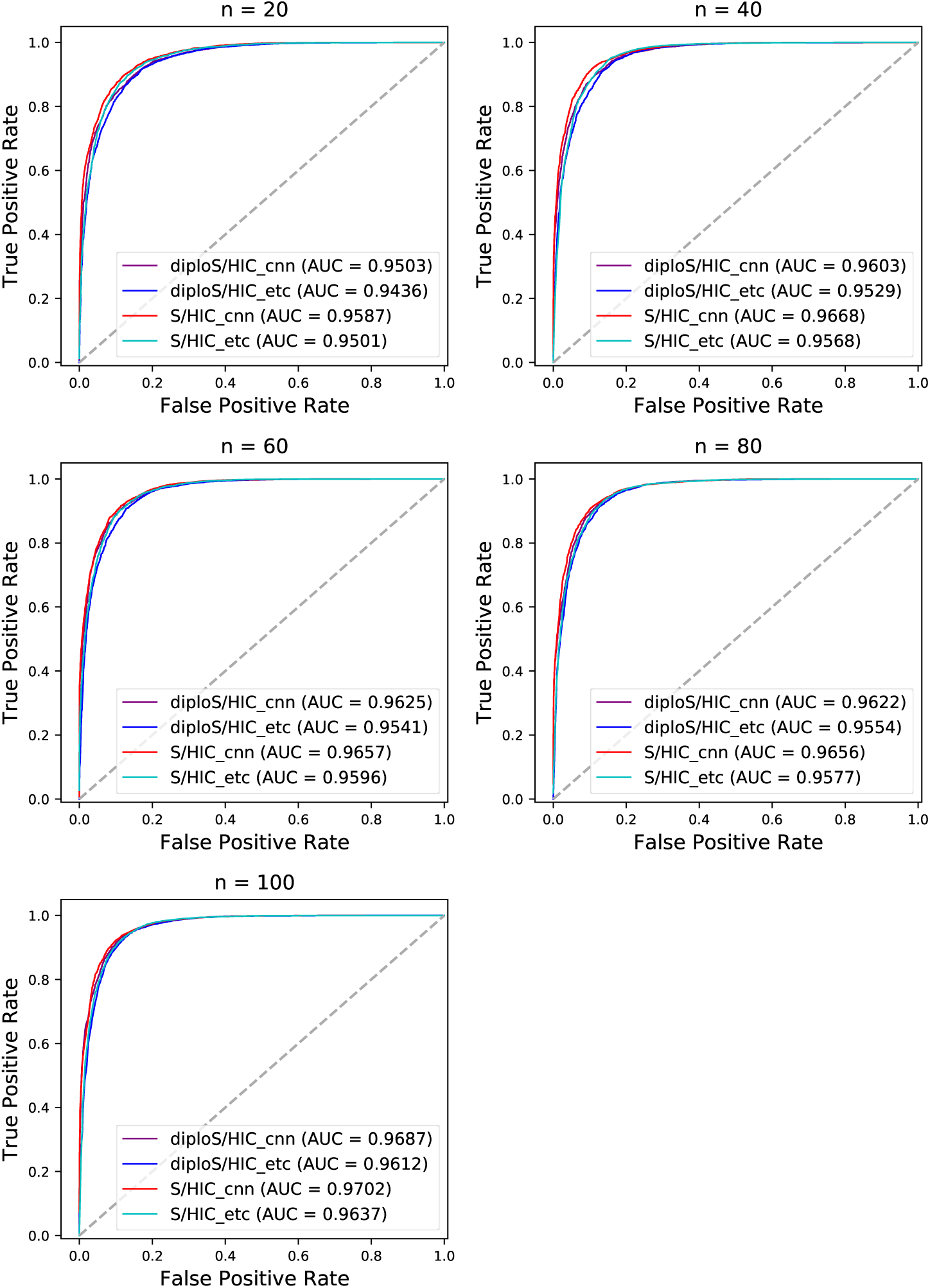
ROC curves showing the true and false positive rates for various classifiers with weak selection. Here we show ROC curves for the binary classification task of Hard and Soft vs linked and neutral regions. ROC curves are shown for both unphased and phased data classifiers as well as for the two machine learning algorithms considered. These simulations were run with the same parameters as those in Figure 6 but with stronger selection, *α ∼ U* (25, 250) Each panel represents a different sample size of haploid chromosomes from *n* = *{*20, 40, 60, 80, 100*}*.

**Figure S6:**
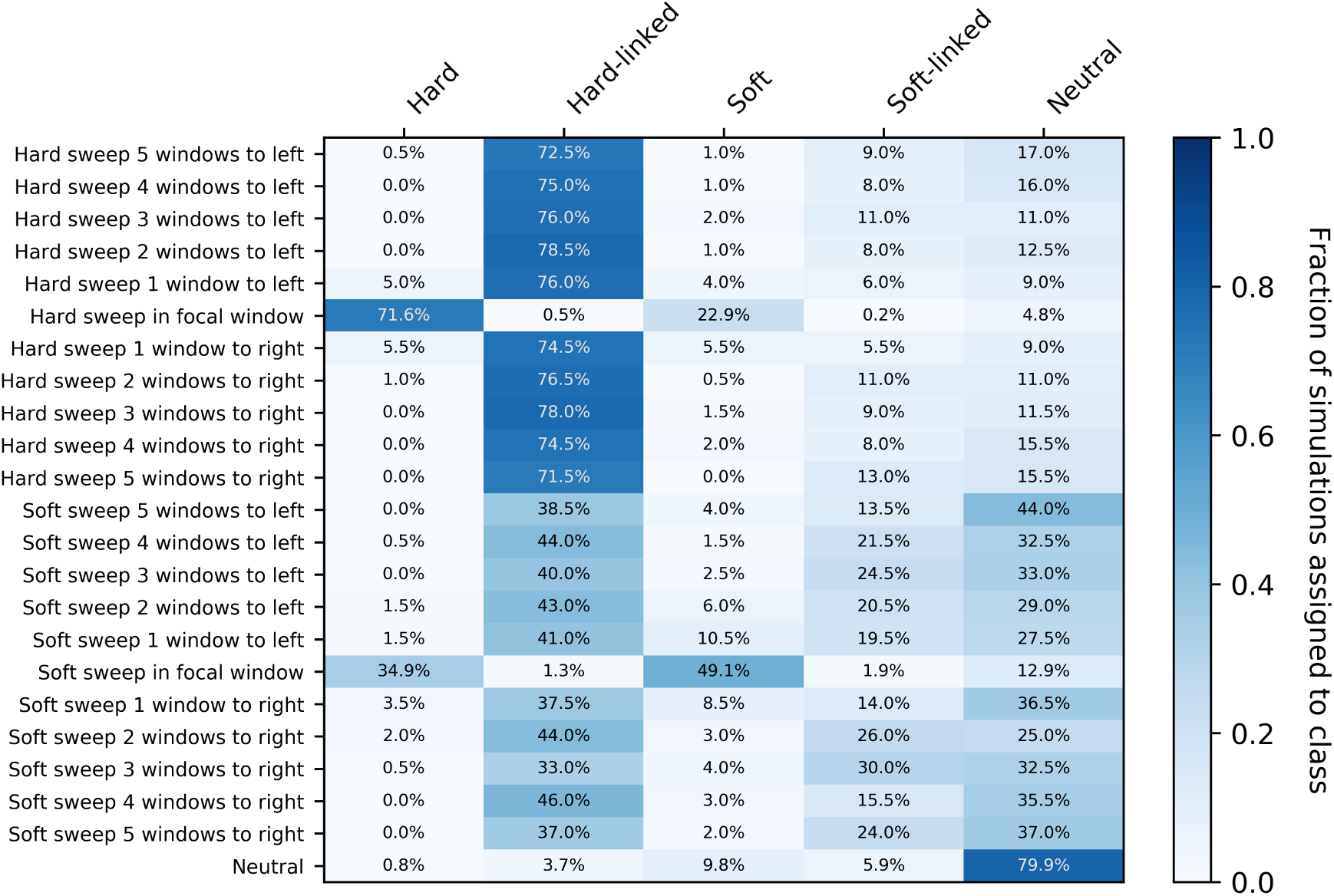
Confusion matrices of subwindows across a recombining chromosome with weak selection. On the y-axis the location of the classified subwindow relative to the sweep is shown while the x-axis shows the predicted class for each sub-window from `diploS/HIC`. These results are from simulations using the same parameter set as in Figure 6 with the exception of selection which is an order of magnitude weaker, *α U* (25, 250), and again using a sample size of *n* = 60 haploid chromosomes.

**Figure S7:**
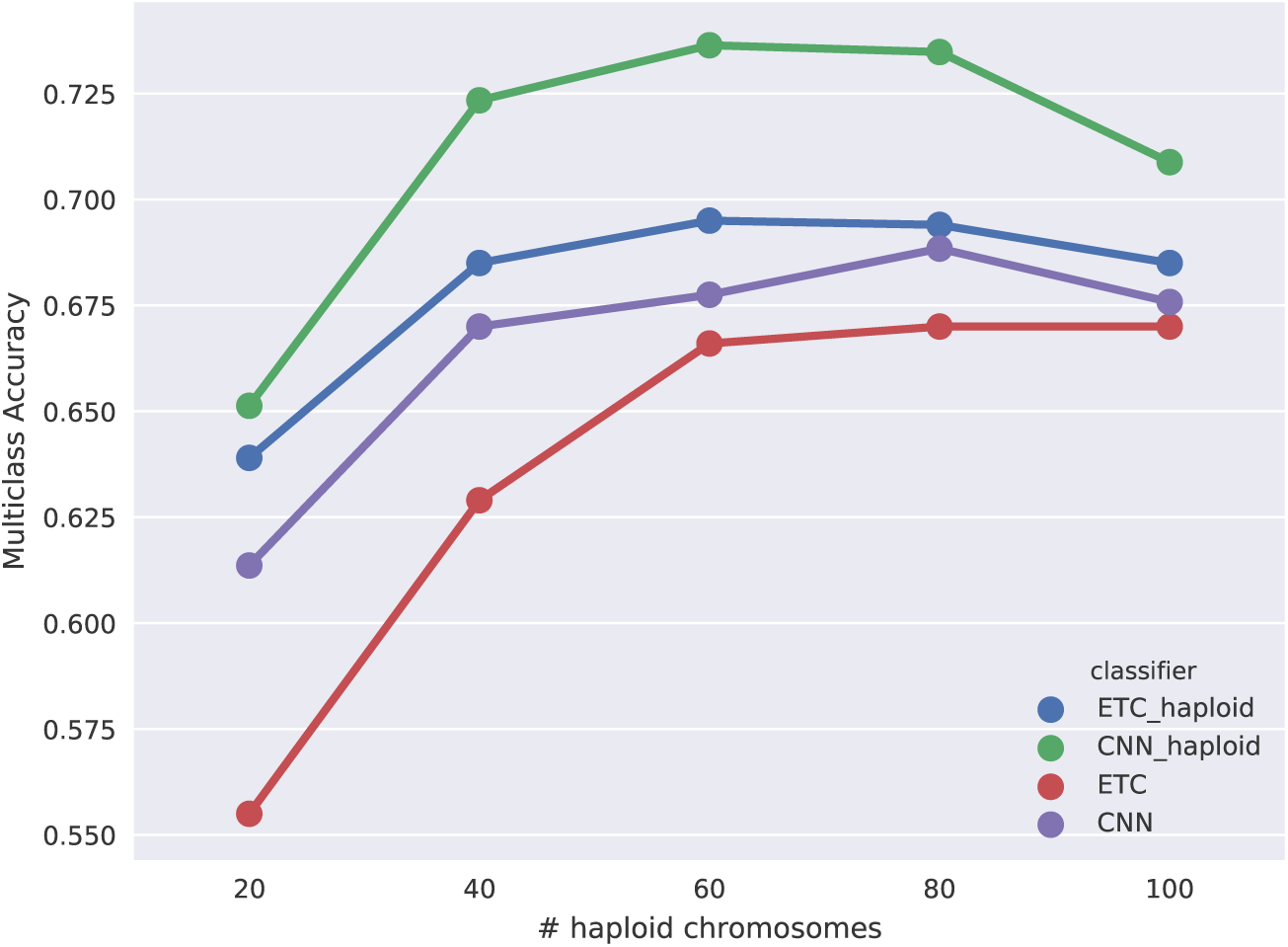
Multiclass classification accuracy for diploS/HIC and related classifiers for *Anopheles* demographic history with strong selection. Here we compare the multiclass classification accuracy of four related classifiers across a variety of sample sizes: the original `S/HIC` classifier (“ETC haploid”), a `S/HIC` classifier which uses a CNN rather than an ETC (“CNN haploid”), our new classifier for unphased data trained with an ETC (“ETC”), and finally our new classifier training with a CNN (“CNN”).

**Figure S8:**
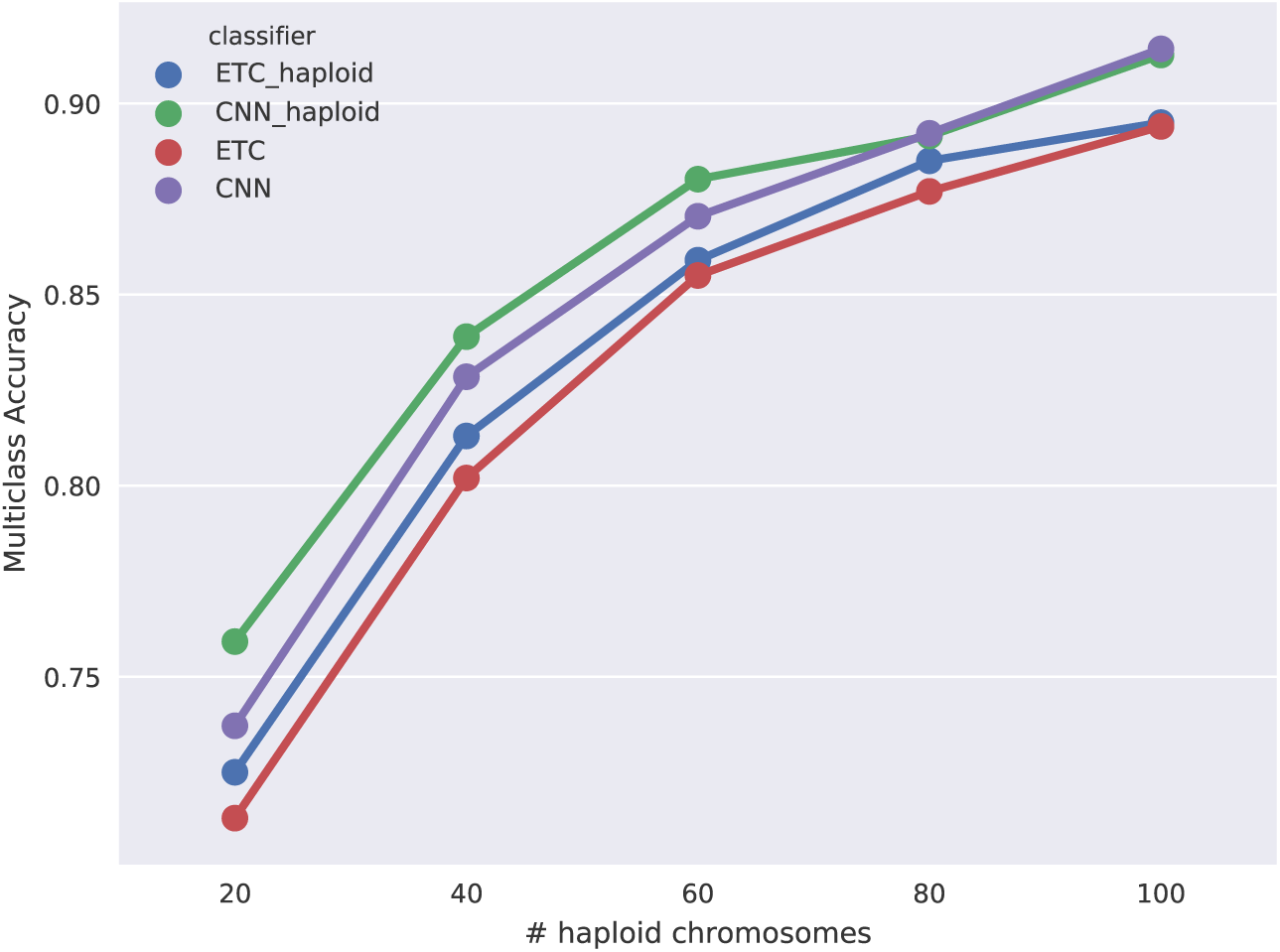
Multiclass classification accuracy for diploS/HIC and related classifiers for *Anopheles* demographic history with weak selection. Here we compare the multiclass classification accuracy of four related classifiers across a variety of sample sizes: the original `S/HIC` classifier (“ETC haploid”), a `S/HIC` classifier which uses a CNN rather than an ETC (“CNN haploid”), our new classifier for unphased data trained with an ETC (“ETC”), and finally our new classifier training with a CNN (“CNN”).

**Figure S9:**
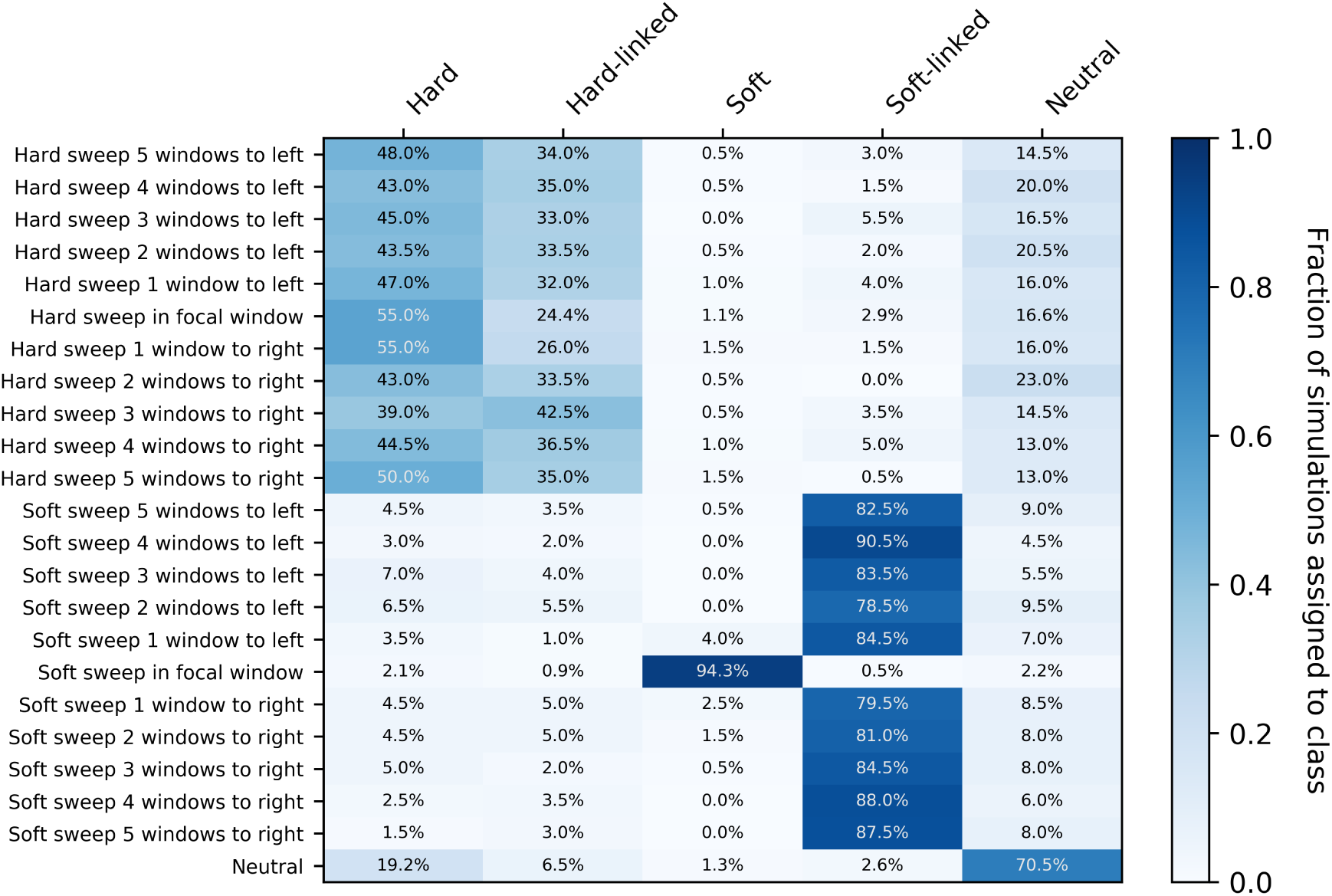
Confusion matrices of diploS/HIC for an *Anopheles gabiae* demographic history with strong selection. On the y-axis the location of the classified subwindow relative to the sweep is shown while the x-axis shows the predicted class for each subwindow from `diploS/HIC`. These results are from simulations using the parameters reflecting the population history of a BFS sample of *Anopheles gambiae*, using a sample size of *n* = 60 haploid chromosomes. Here we have simulated sweeps with very strong selection coefficients *α ∼ U* (2.5 *×* 10^4^, 2.5 *×* 10^5^)

**Figure S10:**
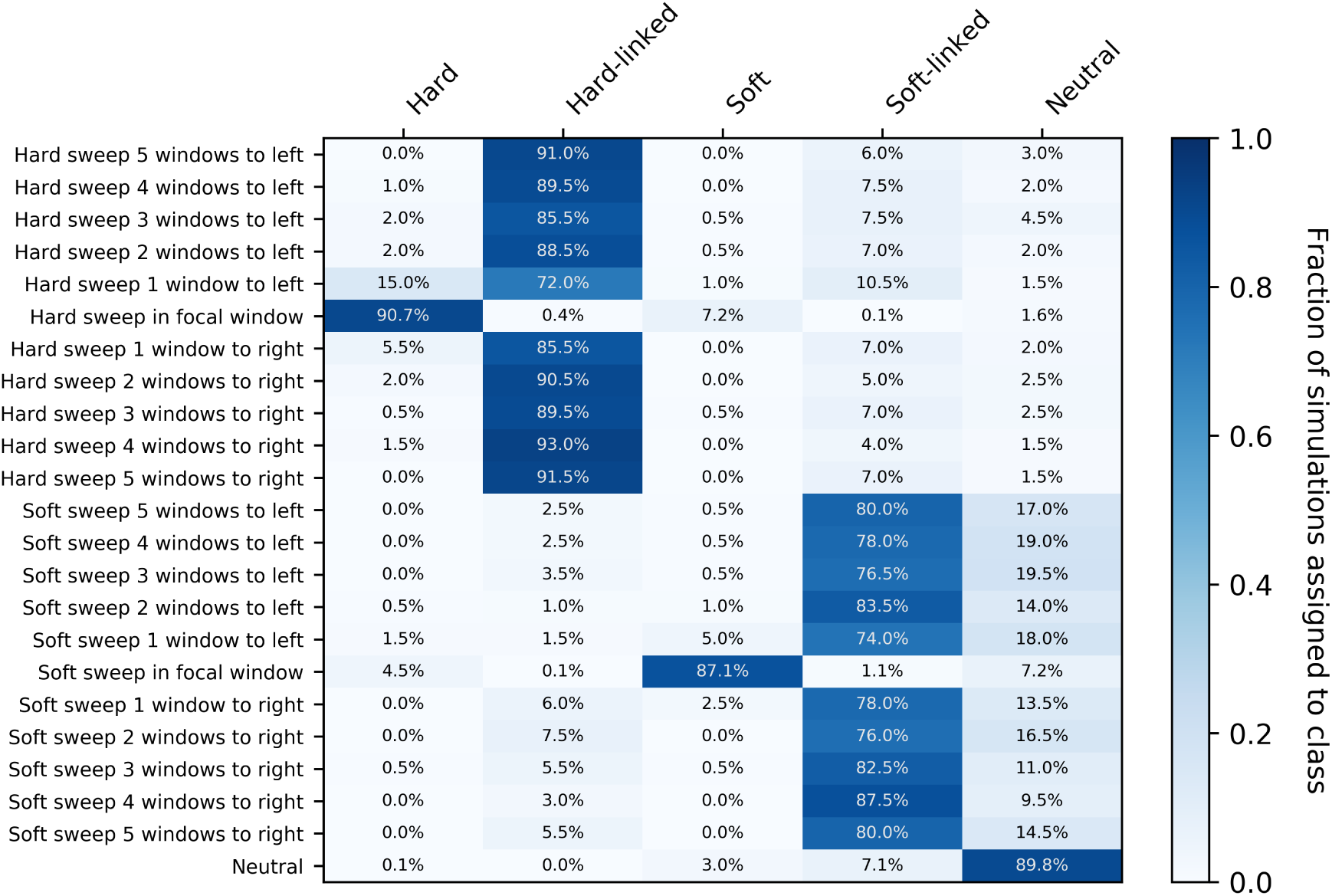
Confusion matrices of diploS/HIC for an *Anopheles gabiae* demographic history with weak selection. On the y-axis the location of the classified subwindow relative to the sweep is shown while the x-axis shows the predicted class for each subwindow from `diploS/HIC`. These results are from simulations using the parameters reflecting the population history of a BFS sample of *Anopheles gambiae*, using a sample size of *n* = 60 haploid chromosomes. Here we have simulated sweeps with weaker selection coefficients,*α ∼ U* (2.5 *×* 10^2^, 2.5 *×* 10^3^)

**Figure S11:**
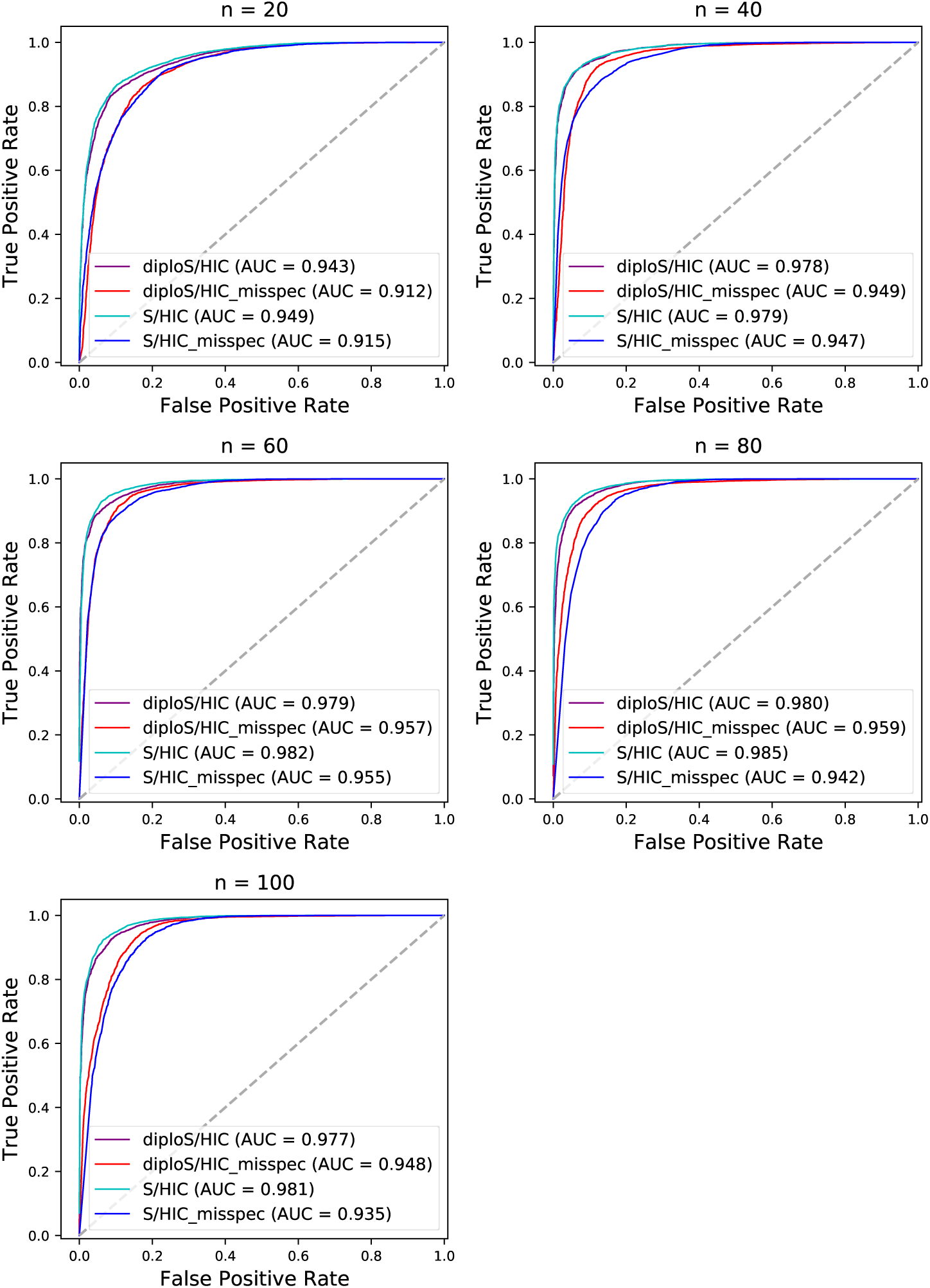
ROC curves showing the effect of model misspecification on performance of diploS/HIC in comparison to S/HIC. Here we compare the performance of `diploS/HIC` and `S/HIC` in the case where the demographic model is terribly mis-specified vs. properly specified. For this example we train using either the BFS *Anopheles* demographic parameters (i.e. Figure 9) or constant population size (i.e. Figure 5) and then test on simulations drawn from BFS parameters. This amounts to catastrophic model misspecification where rather than accounting for the strong population growth in the test set, training ignores the demography completely. As we have shown previously, `S/HIC` is quite robust to such model misspecification, and indeed `diploS/HIC` shares this property.

**Figure S12:**
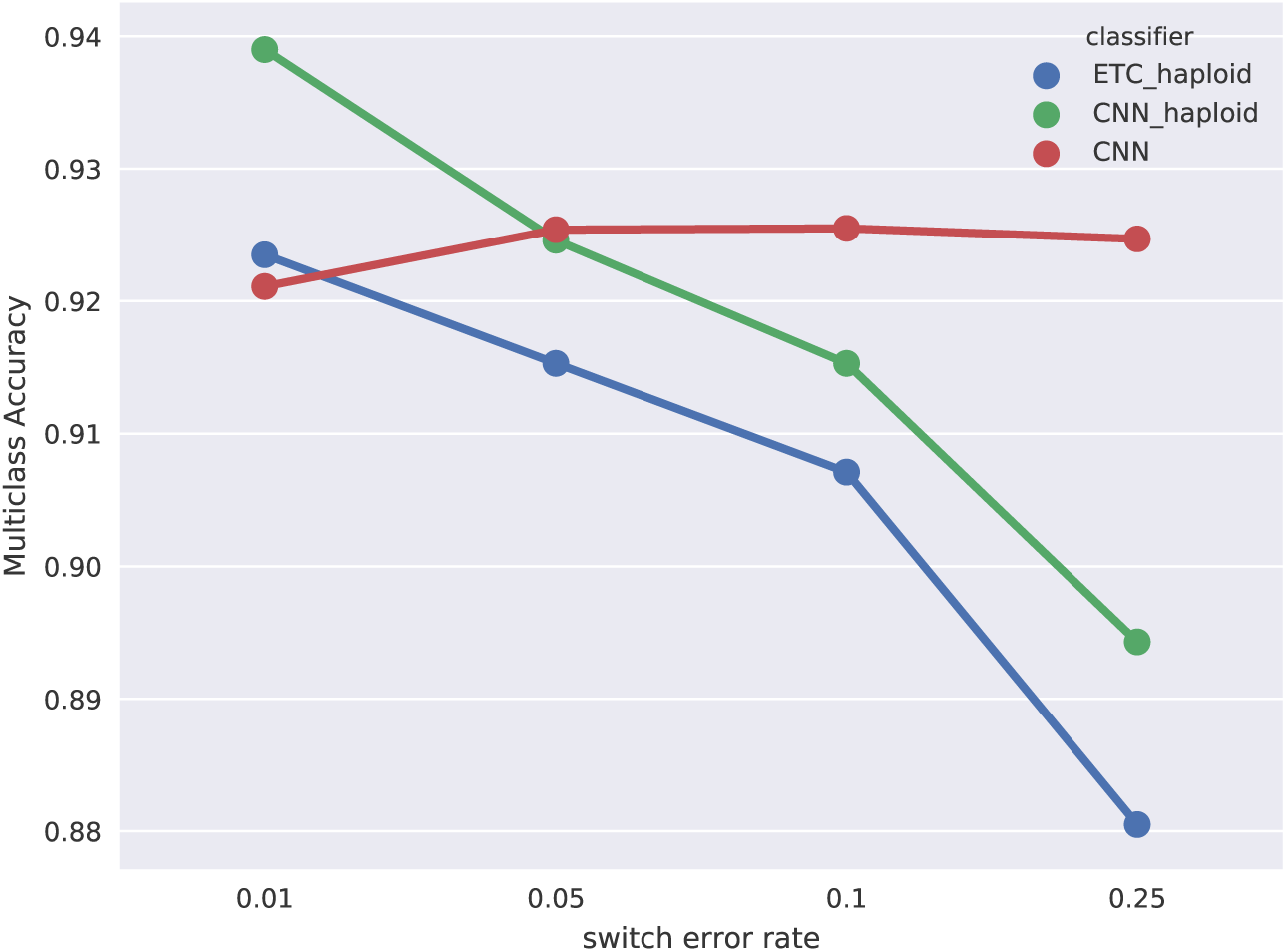
Accuracy of classifiers at various rates of phasing switch error. Here we show multiclass accuracy of three classifers: our original `S/HIC` (labeled ETC haploid), the haploid version of `diploS/HIC` (labeled CNN haploid), and the diploid version of `diploS/HIC` (labeled CNN) as the rate of phasing switch error increases.

## References

Martin Abadi, Ashish Agarwal, Paul Barham, Eugene Brevdo, Zhifeng Chen, Craig Citro, Greg S Corrado, Andy Davis, Jeffrey Dean, Matthieu Devin, et al. Tensorflow: Large-scale machine learning on heterogeneous distributed systems. arXiv preprint arXiv:1603.04467, 2016.

Anopheles gambiae 1000 Genomes Consortium, Data analysis group, Partner working group, Sample collections—Angola:, Burkina Faso:, Cameroon:, Gabon:, Guinea:, Guinea-Bissau:, Kenya:, Uganda:, Crosses:, Sequencing and data production, Web application development, and Project coordination. Genetic diversity of the african malaria vector anopheles gambiae. Nature, 552(7683):96–100, Dec 2017. doi: 10.1038/nature24995.

Jeffrey Chan, Valerio Perrone, Jeffrey P Spence, Paul A Jenkins, Sara Mathieson, and Yun S Song. A likelihood-free inference framework for population genetic data using exchangeable neural networks. arXiv preprint arXiv:1802.06153, 2018.

Francois Chollet et al. Keras, 2015.

Michael DeGiorgio, Christian D Huber, Melissa J Hubisz, Ines Hellmann, and Rasmus Nielsen. Sweepfinder2: increased sensitivity, robustness and flexibility. Bioinformatics, 32(12):1895–7, Jun 2016. doi: 10.1093/bioinformatics/btw051.

J. C. Fay and C. I. Wu. Hitchhiking under positive darwinian selection. Genetics, 155: 1405–1413, 2000.

Nandita R Garud, Philipp W Messer, Erkan O Buzbas, and Dmitri A Petrov. Recent selective sweeps in north american drosophila melanogaster show signatures of soft sweeps. PLoS Genet, 11(2):e1005004, 2015.

Pierre Geurts, Damien Ernst, and Louis Wehenkel. Extremely randomized trees. Machine learning, 63(1):3–42, 2006.

Alex Graves, Abdel-rahman Mohamed, and Geoffrey Hinton. Speech recognition with deep recurrent neural networks. In Acoustics, speech and signal processing (icassp), 2013 ieee international conference on, pages 6645–6649. IEEE, 2013.

Janet Hemingway, Hilary Ranson, Alan Magill, Jan Kolaczinski, Christen Fornadel, John Gimnig, Maureen Coetzee, Frederic Simard, Dabire K Roch, Clement Kerah Hin-zoumbe, et al. Averting a malaria disaster: will insecticide resistance derail malaria control? The Lancet, 387(10029):1785–1788, 2016.

Joachim Hermisson and Pleuni S Pennings. Soft sweeps: molecular population genetics of adaptation from standing genetic variation. Genetics, 169(4):2335–52, 2005. ISSN 0016-6731 (Print).

Jeffrey D Jensen. On the unfounded enthusiasm for soft selective sweeps. Nature Communications, 5:5281, 2014.

Jeffrey D Jensen, Yuseob Kim, Vanessa Bauer DuMont, Charles F Aquadro, and Carlos D Bustamante. Distinguishing between selective sweeps and demography using dna polymorphism data. Genetics, 170(3):1401–1410, 2005.

N. L. Kaplan, R. R. Hudson, and C. H. Langley. The hitchhiking effect revisited. Genetics, 123:887–899, 1989.

John K Kelly. A test of neutrality based on interlocus associations. Genetics, 146(3): 1197–1206, 1997.

Andrew D Kern and Daniel R Schrider. Discoal: flexible coalescent simulations with selection. Bioinformatics, 32(24):3839–3841, Dec 2016. doi: 10.1093/bioinformatics/btw556.

Yuseob Kim and Rasmus Nielsen. Linkage disequilibrium as a signature of selective sweeps. Genetics, 167(3):1513–24, 2004. ISSN 0016-6731 (Print).

Diederik Kingma and Jimmy Ba. Adam: A method for stochastic optimization. arXiv preprint arXiv:1412.6980, 2014.

Alex Krizhevsky and G Hinton. Convolutional deep belief networks on cifar-10. Unpublished manuscript, 40, 2010.

Alex Krizhevsky, Ilya Sutskever, and Geoffrey E Hinton. Imagenet classification with deep convolutional neural networks. In Advances in neural information processing systems, pages 1097–1105, 2012.

Yann LeCun, Fu Jie Huang, and Leon Bottou. Learning methods for generic object recognition with invariance to pose and lighting. In Computer Vision and Pattern Recognition, 2004. CVPR 2004. Proceedings of the 2004 IEEE Computer Society Conference on, volume 2, pages II–104. IEEE, 2004.

Kao Lin, Haipeng Li, Christian Schlotterer, and Andreas Futschik. Distinguishing positive selection from neutral evolution: boosting the performance of summary statistics. Genetics, 187(1):229–44, Jan 2011. doi: 10.1534/genetics.110.122614.

JM. Maynard Smith and J. Haigh. The hitch-hiking effect of a favourable gene. Genetical Research, (23):23–35, 1974.

Philipp W Messer and Dmitri A Petrov. Population genomics of rapid adaptation by soft selective sweeps. Trends Ecol Evol, 28(11):659–69, Nov 2013. doi: 10.1016/j.tree.2013.08.003.

Sara N Mitchell, Daniel J Rigden, Andrew J Dowd, Fang Lu, Craig S Wilding, David Weetman, Samuel Dadzie, Adam M Jenkins, Kimberly Regna, Pelagie Boko, et al. Metabolic and target-site mechanisms combine to confer strong ddt resistance in anopheles gambiae. PLoS One, 9(3):e92662, 2014.

Rasmus Nielsen, Scott Williamson, Yuseob Kim, Melissa J Hubisz, Andrew G Clark, and Carlos Bustamante. Genomic scans for selective sweeps using snp data. Genome Res, 15(11):1566–75, Nov 2005. doi: 10.1101/gr.4252305.

Keiron O’Shea and Ryan Nash. An introduction to convolutional neural networks. arXiv preprint arXiv:1511.08458, 2015.

Pavlos Pavlidis, Jeffrey D Jensen, and Wolfgang Stephan. Searching for footprints of positive selection in whole-genome SNP data from nonequilibrium populations. GENETICS, 185(3):907–922, July 2010.

Ryan Poplin, Dan Newburger, Jojo Dijamco, Nam Nguyen, Dion Loy, Sam S Gross, Cory Y McLean, and Mark A DePristo. Creating a universal snp and small indel variant caller with deep neural networks. BioRxiv, page 092890, 2017.

Marc Pybus, Pierre Luisi, Giovanni Marco Dall’Olio, Manu Uzkudun, Hafid Laayouni, Jaume Bertranpetit, and Johannes Engelken. Hierarchical boosting: a machine-learning framework to detect and classify hard selective sweeps in human populations. Bioinformatics, 31(24):3946–3952, 2015.

Alan R Rogers and Chad Huff. Linkage disequilibrium between loci with unknown phase. Genetics, 182(3):839–844, 2009.

Roy Ronen, Nitin Udpa, Eran Halperin, and Vineet Bafna. Learning natural selection from the site frequency spectrum. Genetics, 195(1):181–93, Sep 2013. doi: 10.1534/genetics.113.152587.

Daniel R Schrider and Andrew D Kern. S/HIC: Robust identification of soft and hard sweeps using machine learning. PLoS Genet, 12(3):e1005928, Mar 2016. doi: 10.1371/journal.pgen.1005928.

Daniel R Schrider and Andrew D Kern. Soft sweeps are the dominant mode of adaptation in the human genome. Mol Biol Evol, 34(8):1863–1877, Aug 2017. doi: 10.1093/molbev/msx154.

Daniel R Schrider and Andrew D Kern. Supervised machine learning for population genetics: A new paradigm. Trends Genet, Jan 2018. doi: 10.1016/j.tig.2017.12.005.

Daniel R Schrider, Fabio K Mendes, Matthew W Hahn, and Andrew D Kern. Soft shoulders ahead: spurious signatures of soft and partial selective sweeps result from linked hard sweeps. Genetics, 200(1):267–84, May 2015. doi: 10.1534/genetics.115.174912.

Daniel R Schrider, Alexander G Shanku, and Andrew D Kern. Effects of linked selective sweeps on demographic inference and model selection. Genetics, 204(3):1207–1223, Nov 2016. doi: 10.1534/genetics.116.190223.

Sara Sheehan and Yun S Song. Deep learning for population genetic inference. PLoS Comput Biol, 12(3):e1004845, Mar 2016. doi: 10.1371/journal.pcbi.1004845.

Katy L Simonsen, Gary A Churchill, and Charles F Aquadro. Properties of statistical tests of neutrality for dna polymorphism data. Genetics, 141(1):413–429, 1995.

Ilya Sutskever, Oriol Vinyals, and Quoc V Le. Sequence to sequence learning with neural networks. In Advances in neural information processing systems, pages 3104–3112, 2014.

Fumio Tajima. Evolutionary relationship of dna sequences in finite populations. Genetics, 105:437–460, 1983.

Fumio Tajima. Statistical method for testing the neutral mutation hypothesis by dna polymorphism. Genetics, 123:585–595, 1989.

G. A. Watterson. On the number of segregating sites in genetical models without recombination. Theoretical Population Biology, 7:256–276, 1975.

Fisher Yu and Vladlen Koltun. Multi-scale context aggregation by dilated convolutions. arXiv preprint arXiv:1511.07122, 2015.

